# FlowTransOP: Distributional Translation of Omics Signatures via Constrained Deep Flow Matching

**DOI:** 10.64898/2026.05.27.728305

**Authors:** Nikolaos Meimetis, Trong Nghia Hoang, Sara Magliacane, Douglas A. Lauffenburger

## Abstract

Observations from pre-clinical models rarely generalize to human patients, leading to many failures in clinical trials. Most existing methods cannot handle domains with non-overlapping features and no paired samples. Here, we developed FlowTransOP to translate biological observations across such domains without requiring 1-to-1 feature mappings and paired data, while providing a guideline for model selection across four translational regimes. We use flow matching to align full domain distributions in a pre-aligned latent space, with a structural regularization term that keeps similar conditions proximate after transformation. FlowTransOP remains competitive with gold-standard approaches requiring paired samples, but outperforms them when pairs become scarce (<35 pairs) or when cross-domain features are only moderately correlated (r<=0.58). Overall, FlowTransOP can translate perturbations between pre-clinical models and patients when direct correspondences are unavailable, enabling reliable therapeutic inference. As a proof-of-concept, we trained a foundational mouse-human transcriptomic map on ARCHS4 and applied it to liver disease predictions.

## 1. Introduction

Translation between different domains and contexts is a fundamental challenge governing many diverse fields, ranging from language translation to the translation of observations between disparate biological systems^1–5^. For the latter, biologists and biomedical researchers often develop and test hypotheses in model organisms (e.g., mice, zebrafish) or in vitro cell cultures, to gain insights concerning the operation of biological processes. However, biology is highly context-specific, and often the observations made in one system do not generalize readily across biological systems (i.e., cells, species, platforms)^6–8^. Importantly, for therapeutics development, there is a critical need to translate observations from pre-clinical experimental models to human patients, as many drugs and vaccines that appear promising in models ultimately fail in the clinic^9,10^. This is due to difficulty in translating the insights gained from these biological models^5,10^, arising from divergence of molecular process effects in human versus animal models^11–15^. This need is also reflected in recent industry efforts to interpret mouse perturbation experiments through human-centered biological readouts, highlighting a growing recognition that scalable preclinical perturbation data are most useful when they can be projected into a human biological coordinate system^16^.

Translating between domains that have non-overlapping feature spaces and limited (or even zero) paired samples is a fundamentally hard problem that intersects biology and machine learning. Basically, when sharing the same measured biological species, this can be interpreted as a homogeneous domain adaptation problem, while inter-species translation with vastly different features is a heterogeneous domain adaptation problem^17,18^. In the biological context, non-overlapping features mean that, for instance, one dataset might be measured in terms of some genes or immune markers, while another dataset uses an entirely different set, where there is no direct feature correspondence. Most existing methods cannot handle this scenario without additional information.

A spectrum of approaches has been proposed to translate observations between different contexts with homogeneous domains. FIT^19^ and DeepCellState^20^ propose a linear and a non-linear (autoencoder-based) approach, respectively, to directly translate omics profiles from one biological system to another, e.g., drug-modulated gene expression from mouse to human. TransCompR^21^, PRECISE^22^, and TRANSACT^23^ align data coming from different biological contexts into a shared latent space, enabling results such as: direct transfer of labels from one system to another; downstream predictions for a target system given new observations on a reference system; or identifying signatures in the reference system that are the most predictive for the target system. However, all these approaches require a 1-1 mapping between the features of the input spaces or suffer a significant performance drop for heterogeneous domains. Even approaches that leverage deep generative models assume some overlap; scGEN^24^ and CPA^25^ can predict single-cell perturbation responses in an unobserved cell type or in a different species by performing vector arithmetic in a latent gene expression space, but still require mapping orthologous genes between species.

For highly heterogeneous inter-species domains typical in biological translation, multimodal alignment using some kind of sample pairing can be performed to bypass the requirement of 1-1 mapping between features. For example, SATL^17^ enables downstream labeling (such as cell annotation) using common labels between biological systems, to construct a shared latent space, building on Cross-Domain Structural Preserving Projection (CDSPP)^26^. SATURN^27^ proposes a different route to cross-species alignment in the single-cell setting, where it couples scRNA-seq counts with protein language model embeddings to group genes into shared macrogenes, enabling integration and label transfer across evolutionarily remote species without restricting analysis to homologs. Even more recently, large cross-species transcriptomic foundation models such as TranscriptFormer have shown that single-cell representations can be learned across broad evolutionary scales, spanning many species, tissues, cell types, and disease states^28^. However, TranscriptFormer is not designed to translate specific source samples into corresponding target domain omics profiles, and does not directly address translating unpaired perturbation-level omics (of any modality) distributions between preclinical models and human disease states when feature spaces do not overlap. AutoTransOP^9^ employs autoencoders to translate between biological contexts exhibiting completely different features by using a relatively small set of condition pairs. This pairing is required to build a global latent space and force “comparable” conditions to be close to each other, regardless of the system of origin. Similarly, CLIP and CLAP use contrastive alignment across visual and audio modalities but require tens of millions of labeled pairs^29,30^. STRUCTURE introduced a modular alignment strategy that dropped the pairing requirement to a few thousand pairs^31^. AutoTransOP at present reports the lowest number of required paired conditions and sample size, with the greatest flexibility in downstream tasks and feature similarity. However, AutoTransOP and the other techniques still require paired data, a situation often unavailable in pre-clinical drug development, which can be cast as a distributional translation task where the patients’ and pre-clinical models’ (e.g., mice) distributions are available without detailed pairing of conditions. No prior solution simultaneously handles completely distinct feature sets and entirely unpaired training data.

Generative AI methods based on diffusion processes and flow matching offer an alternative paradigm for unpaired domain translation, promising improved biological translation. An initial transfer between patient and pre-clinical (e.g., animals, in vitro models, etc.) distributions is still required before going on a clinical trial, even in the absence of pairings, so such methods can aid in modeling the transition between these distributions. For instance, Brownian Bridge Diffusion Models (BBDM) treat image-to-image translation as learning a continuous stochastic bridge between source and target domains rather than as direct conditional generation^2^. Similarly, the emerging Flow Matching approaches seek to learn a smooth mapping (e.g., a velocity field) that transforms a source distribution into a target distribution by matching probability flows between them^32^. Learning such probabilistic bridges for transforming distributions from a source to a target distribution has already been successful in offline black-box optimization tasks, using ROOT^33^. Thus, utilizing flow matching and diffusion techniques with appropriate constraints for preserving structure in the data could enable, to some extent, a pair-free and homologue-free translation between biological systems.

In this work, we introduce a new method (termed FlowTransOP) to translate biological observations across disparate domains without requiring one-to-one feature mappings and without paired data. We present a framework for determining the optimal translation approach across four difficulty regimes: (i) fully paired domains with shared features, (ii) limited pairing with shared features, (iii) completely unpaired domains with correlated features, and (iv) completely unpaired domains with distinct or uncorrelated features — a scenario frequently encountered in therapeutic development. Within this context, we highlight where FlowTransOP provides distinct advantages over established gold-standard methods like AutoTransOP. Specifically, we leverage a flow matching generative model to learn a mapping between two domains (e.g., cell line A and cell line B) by directly aligning their data distributions. Intuitively, instead of assuming that we operate on the same feature space or require paired data, we learn a transformation that carries the entire feature profile by matching the underlying probability flows. To improve stability, we incorporate a pre-aligned latent space as an initial guide. We use TRANSACT to obtain a coarse shared embedding of the two domains (even if it is inaccurate at the beginning), but any pre-alignment technique can be used. Then we use a flow-based model to perform conditional generation from one domain to the other along that embedded manifold. This constitutes FlowTransOP, a fully unpaired generative translation framework that leverages flow matching to align the entire distributions of distinct domains while enforcing structural consistency through a pre-aligned latent space. By systematically evaluating FlowTransOP across the four regimes of translational difficulty, we demonstrate that it outperforms naive and random baselines and remains competitive with contrastive approaches when sufficient pairing exists. When pairs are scarce or absent, or when features are uncorrelated, FlowTransOP uniquely maintains meaningful predictive performance, whereas all other methods fail. Beyond introducing this method, our work defines a benchmark and provides guidelines for selecting between flow-based, latent decoders, and contrastive methods depending on feature similarity and pairing availability. Finally, we demonstrate the therapeutic relevance of this framework by training a foundational mouse-human map from the ARCHS4 dataset^34^ and using it to translate mouse liver perturbation signatures into human MASH-associated disease scores.

## 2. Results

### 2.1. Translational tasks span a spectrum of difficulty from paired to unpaired domains

A hierarchy of translation difficulty can be identified, based on two factors: i) the similarity and overlap of the features of the input domains, and ii) the availability of paired samples between the two domains (Figure 1). Having highly dissimilar molecular features as input, such as nonhomologous genes or distinct adaptive immune system components in biological omics studies, is the toughest case one can encounter. Meanwhile, having the same homogenous feature space, such as the same genes or genes that are at least very correlated, is the easiest case. Approaches like AutoTransOP perform exceptionally well even in cases where the input features are completely different between two biological systems; however, performance there drops significantly when the number of paired samples between the two domains becomes low^9^. Having few or even zero pairs is unfortunately common in drug development studies, where pharmaceutical companies may possess plenteous data for a specific drug tested on pre-clinical and animal models, but only disparate public datasets from human patients with the disease, but untreated (since the envisioned drug is under development). AutoTransOP cannot be trained in the absence of paired conditions, and other approaches utilizing some approximate alignment have low performance in cases with highly dissimilar domains. Hence, we develop here a new flow matching-based approach, called FlowTransOP (section 2.2, Figure 2a), to simultaneously address completely distinct feature sets and entirely unpaired training data. For assessment of this method, we define four regimes of difficulty (Figure 1): (i) fully paired domains with shared features, (ii) limited pairing with shared features, (iii) completely unpaired with correlated features, and (iv) completely unpaired with distinct or uncorrelated features (the hardest possible regime but the most realistic one for therapeutic development). We use the publicly available L1000 dataset^35^, firstly, as it was originally pre-processed and divided into train and test splits in the AutoTransOP framework^9^, to ensure fairness in comparing with the state-of-the-art. However, we implement our own evaluation regimes on it to ultimately create a benchmark dataset for translation tasks of different difficulties (Figure 1). This strategy allows us to systematically identify when simple latent space methods suffice and when more sophisticated approaches, such as flow matching, are necessary.

**Figure 1:**
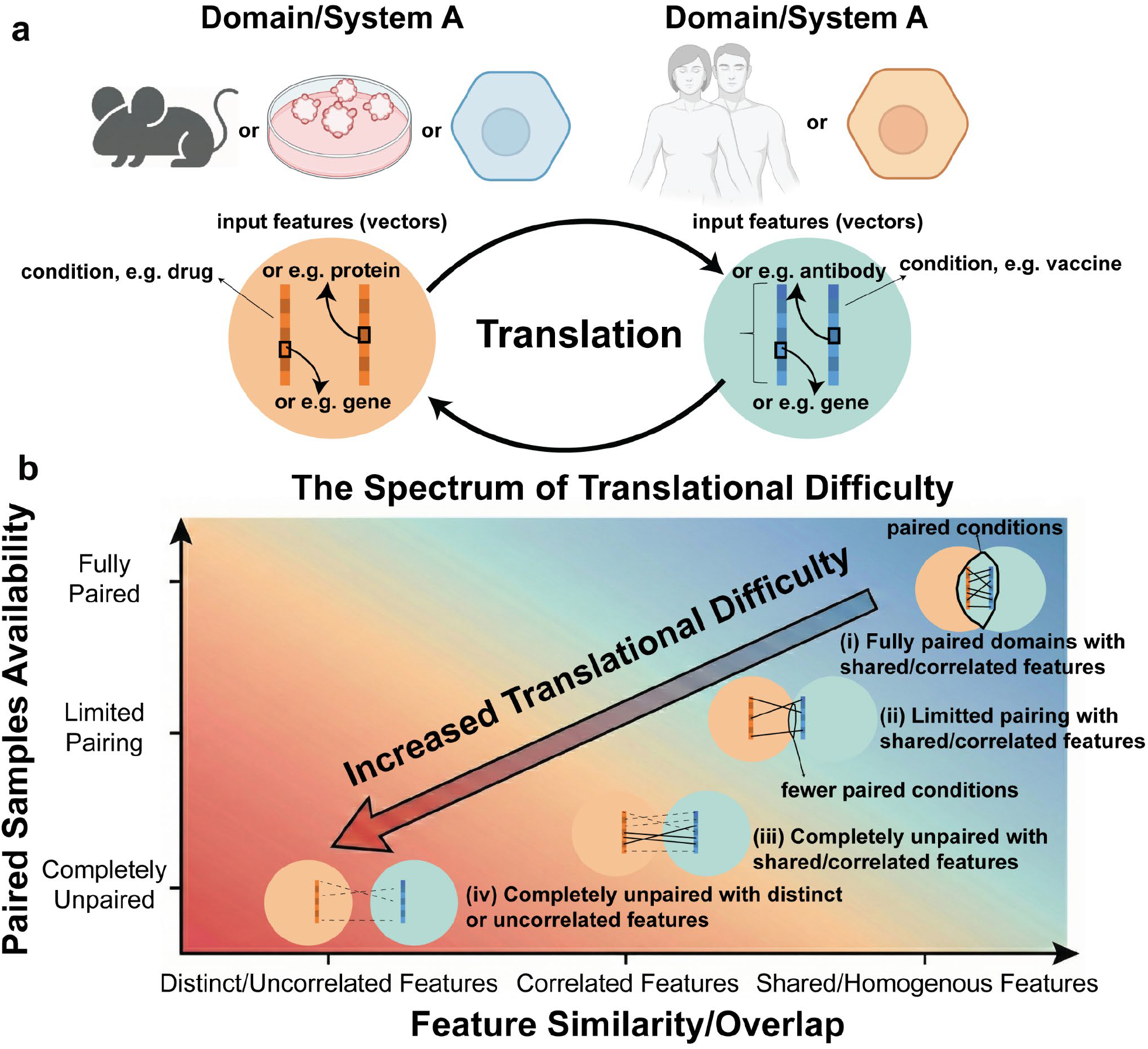
The Spectrum of Translational Difficulty. There are four regimes of difficulty, where different approaches are expected to generally perform best. These are: (i) fully paired domains with shared features, (ii) limited pairing with shared features, (iii) completely unpaired with correlated features, and (iv) completely unpaired with distinct or uncorrelated features.

**Figure 2:**
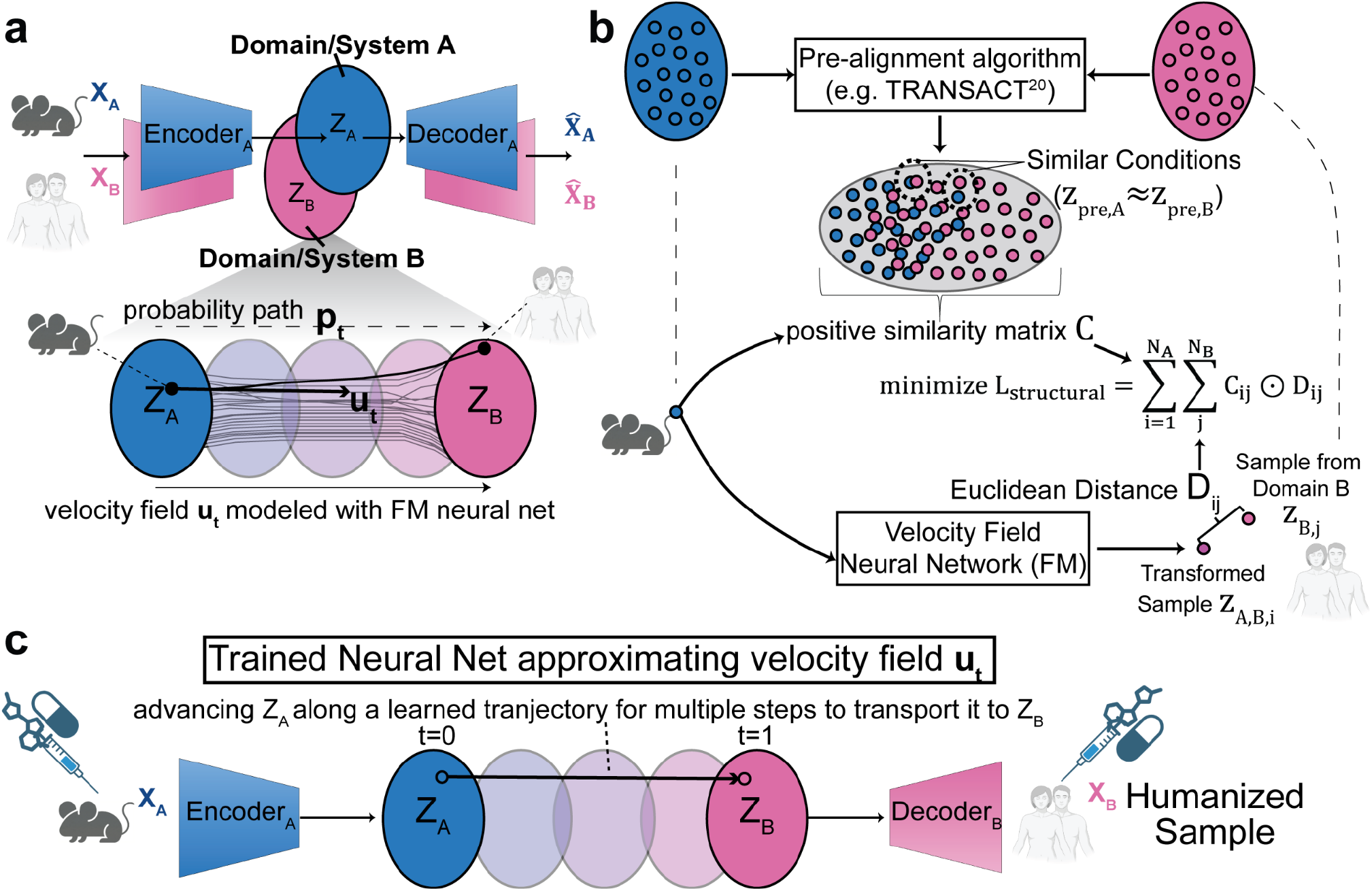
Schematic of the unpaired deep learning approach for generalized translation, which utilizes constrained Flow Matching (FM). **a)** Pre-training separate autoencoders for each domain to embed samples into latent distributions of the same dimensionality. Then, an FM neural network is trained to transform data from a source domain to a target domain. **b)** Domains are pre-aligned to identify approximate similarity of samples and force structure during training the flow matching model, so that approximately similar conditions are close to each other after the transformation is applied. **c)** Inference step of the model. After having trained each component, a new unseen perturbation in a source domain (e.g., a drug tested on mice) is translated to a target domain (e.g., humans in the clinic).

### 2.2. FlowTransOP: A Flow-Matching Framework for Unpaired Translation

For the case of unpaired datasets, we can treat unpaired biological translation as a distributional alignment problem, rather than a sample-to-sample mapping (Figure 1, Figure 2a). This means that we wish to transform a source distribution (e.g., cell line A, or species A, etc.) to a target distribution (e.g., cell line B, or species B, etc.). Flow Matching (FM) is suitable for this task as it uses simple regression to train a neural net to learn a velocity field (*u*_*t*_) that can be later used to build a probability path (*p*_*t*_) from a source distribution to a target distribution^32^ (see Materials and Methods for details). During inference, we may draw a sample from the source distribution and transform it into the target distribution by solving the Ordinary Differential Equation (ODE) determined by the velocity field. To use this with different input domains, we pre-train and fix a separate autoencoder for each domain (here, biological system, i.e., different cell lines in the benchmark dataset from the L1000 dataset^35^) to embed our input representation into latent representations of the same dimensionality, but clearly representing different distributions. Then, we train two different velocity fields to transform data from one latent space to another. In practice, FlowTransOP samples latent source and target points within each minibatch, constructs intermediate states along a linear conditional optimal-transport path, and trains the neural velocity field to predict the displacement between the two latent states. Thus, the core flow matching objective is a supervised regression problem in latent space, while the learned velocity field is later integrated to transport unseen source samples into the target latent distribution. However, FM by itself does not guarantee that the transformation from one space to another results in similar conditions being close to each other, e.g., the same drug in cell line A might not be mapped to a latent representation close to the vicinity of its latent representation in cell line B, which would be required by the decoder of cell line B to correctly translate at the end the input features. In fact, training an FM model with no constraint at all in transforming distribution results basically in performance similar to simply predicting that a drug has the same effect on the two cell lines (Supplementary Figure S1), which is simply not true.

To incorporate structural guidance, we utilize a pre-aligned latent space to provide an initial guide (even if somewhat inaccurate), improving the stability and accuracy of the conditional generation (Figure 2b). We reasoned that even if the pre-aligned space is not sufficiently accurate, it may be adequate for providing constraints on the velocity field neural network. Here, we utilized TRANSACT^23^, which identifies common processes between domains and embeds them into a shared mathematical space (see Materials and Methods). A user may utilize other algorithms for pre-alignment, but each technique contains different levels of information influencing downstream capability. For example, using TRANSACT^23^ pre-aligned spaces with FM significantly outperforms using STRUCTURE^31^ with FM (Supplementary Figure S2). After pre-alignment, a regularization term is added to the loss function during training, where similar conditions in the pre-aligned space are forced to be close after transforming the source to the target distribution (Figure 2b), via Euclidean distance minimization. We also evaluated (section 2.3) whether simply training decoders to map back to the original input spaces is sufficient to enable unpaired translation.

Once training is complete, translation is enabled by utilizing the autoencoders and the learned velocity field. Specifically, consider translating a perturbation from system A to system B (Figure 2c): during inference time, the input features of system A (under some condition) are embedded into their own latent space via their corresponding encoder A. Then the trained neural network approximating the velocity field is used to transform that latent representation into another representation in the latent space of system B. Finally, the corresponding decoder B of the other system is used to translate back into the input features of system B.

Finally, tested the potential of training simple decoders in pre-aligned space to enable translation. The pre-alignment serves to create an approximately global space for the two domains. We reasoned that even if that space is approximate in the case of different features, some structural information is contained, so training decoders to decode projected data from each domain back to their corresponding input space could enable translation, at least in easier regimes.

### 2.3. Performance in the Ideal Case: Benchmarking Simple Translation

First, it is essential to validate that our constrained FM approach, FlowTransOP, can outperform naive baselines in the easiest of the regimes, i.e., fully paired domains with shared features. For a naive baseline, we use direct translation of a perturbation from one cell line to another, assuming that the observed drug-induced gene expression in cell line A is the same in cell line B. Additionally, we train FM-based models using shuffled features to establish a random baseline for the approach itself. In this regime, contrastive learning-based approaches like AutoTransOP are expected to be the best option and to set an upper bound on achievable performance. However, FM should at least be better than the naive and random baselines. Finally, we also validated the potential of training simple decoders in the pre-aligned space to translate between domains.

To validate each approach, we utilized a 5-fold cross-validation strategy (only for testing, not hyperparameter tuning). For evaluating translation performance, we use samples derived from the same drug, tested on the same dose and time duration in 2 different cell lines, not seen by any model component during training. This was implemented across 8 cell line pairs from the L1000 dataset, where AutoTransOP’s performance was also calculated^9^. We observed that indeed AutoTransOP serves as an upper bound baseline of performance since, when there is a sufficient number of pairs, it outperforms FlowTransOP (Figure 3a), as well as just using decoders in the pre-aligned space (Figure 3c). FlowTransOP always outperforms the random baseline, while it is better than the naive baseline of direct translation for most cases (Figure 3a), establishing that FM can indeed be used to learn meaningful cross-domain mappings, even in the complete absence of paired data information. In this regime, where cell lines share the same features, pre-alignment works so well that simply training decoders in that pre-aligned space is enough to achieve high performance, better than using a more complex FM-based approach (Figure 3b). However, using only pre-alignment performs significantly worse than FlowTransOP (in fact, it performs the same as random models) in the hardest regimes with completely different uncorrelated features (Supplementary Figure S3, Figure 4, more details in section 2.5). Moreover, when utilizing actual pairs to constrain FM (instead of similarity in the pre-aligned space), FlowTransOP generalizes the same as the decoders-only approach, but it fits the data better in training (Supplementary Figure S4). By employing a hybrid strategy that combines available sample pairs with similarity constraints in a pre-aligned space, as detailed in the Methods section, we achieve performance levels comparable to high-pairing scenarios while maintaining greater robustness when pairs are limited (Figure 3d). This approach serves as an effective middle ground, offering a strategic trade-off between the contrastive-based AutoTransOP and the entirely unpaired FlowTransOP framework. Finally, as noted earlier, different pre-alignment methods contain different amounts of information content (Supplementary Figure S2); for the chosen TRANSACT approach, we need to build the consensus space using both domains (Supplementary Figure S5). Which domain is used as a source or target does not matter for the pre-alignment, while for a sufficient batch size, whether the pre-alignment contains paired samples or not has no significant effect (Supplementary Figure S5).

**Figure 3:**
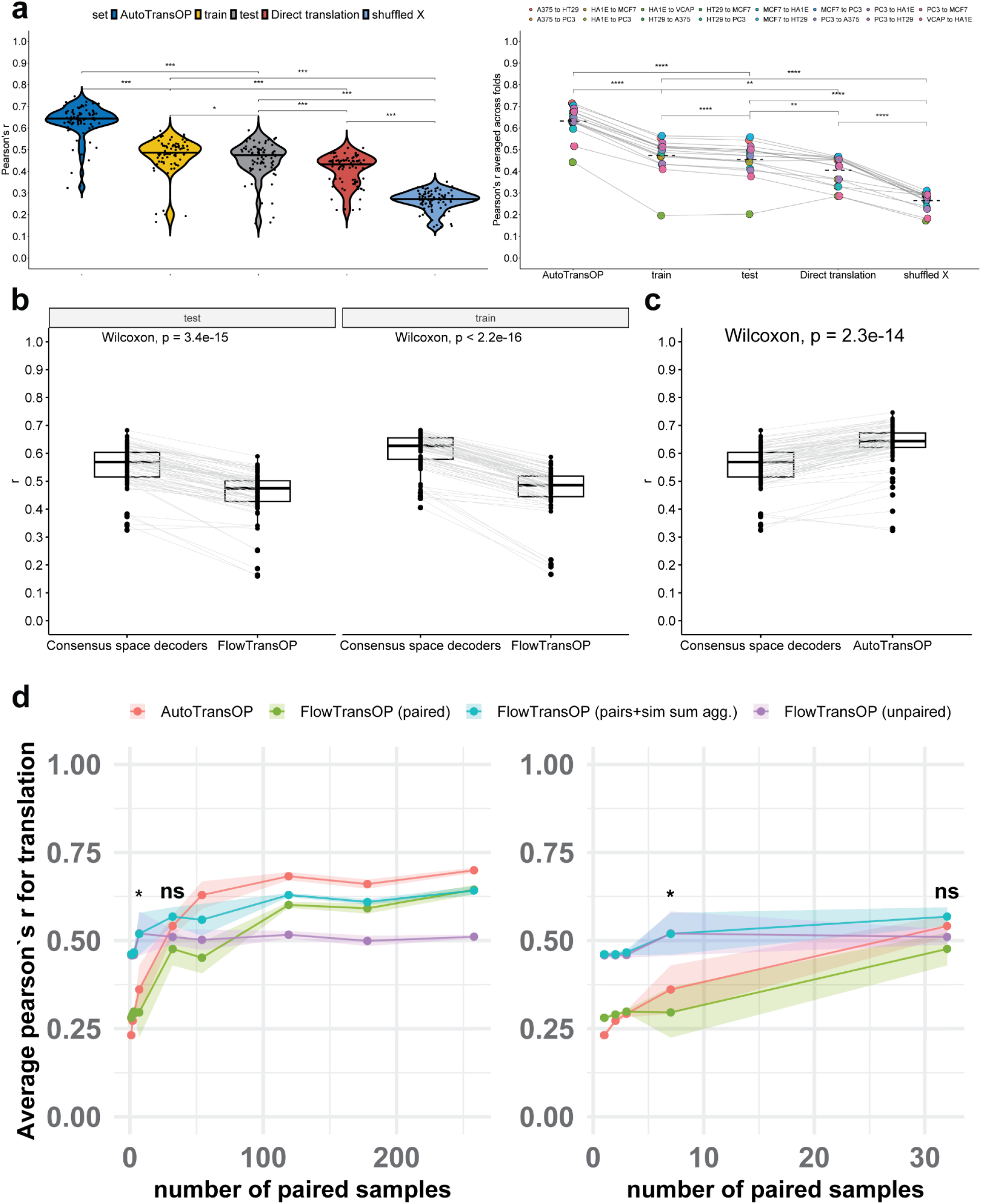
Validation of FlowTransOP against naive, random, and upper-bound baselines in the easiest translation regime of domains with shared features and paired samples. **a)** Translation performance across L1000 cell-line pairs in the fully paired, shared-feature setting, measured as Pearson’s correlation (r) between predicted and observed gene expression profiles under 5-fold CV. Methods include AutoTransOP, FlowTransOP evaluated on training (train) and held-out (test) folds, a direct translation baseline that assumes the perturbation response is the same between source and target cell lines, and a shuffled X random baseline, where an FM model is trained on shuffled genes. **Left**, violin plots show fold-level performance across all directed translation tasks. Statistical annotations correspond to prespecified pairwise contrasts from a linear mixed-effects model fit to Fisher z-transformed correlations, with method as a fixed effect and translation task and fold as random intercepts. P-values were adjusted using the Bonferroni correction. **Right**, each point denotes the mean performance of one directed translation task averaged across folds, with gray lines connecting the same task across methods. Significance was assessed using paired Wilcoxon signed-rank tests. **b)** Comparison between simple decoders trained in the pre-aligned consensus space (using TRANSACT^23^) and FlowTransOP, shown separately for held-out test folds (**left**) and training folds (**right**). Each line connects matched translation tasks, allowing direct within-task comparison. P-values were computed using paired Wilcoxon signed-rank tests. **c)** Comparison of consensus space decoders with AutoTransOP on held-out test data. Each line connects the same directed translation task across methods. Statistical significance was assessed using a paired Wilcoxon signed-rank test. **d)** Comparison of the performance of AutoTransOP and FlowTransOP as the number of paired training conditions is progressively reduced in the low-pair benchmark. For FlowTransOP, we show training different regimes of utilizing fully unpaired training, paired training, and a mix of known pairs and similarity in the pre-aligned space. Shown are the mean held-out translation accuracies (Pearson’s r) across 5-fold CV splits. Shaded ribbons denote a deviation of 1 standard error of the mean (S.E.). The **left** panel shows performance across a wide range of numbers of paired samples, while the **right** panel zooms in on the lowest number of available pairs. Significance labels indicate paired t-tests on Fisher z-transformed fold-level correlations comparing AutoTransOP and unpaired FlowTransOP at the marked paired sample counts.

**Figure 4.**
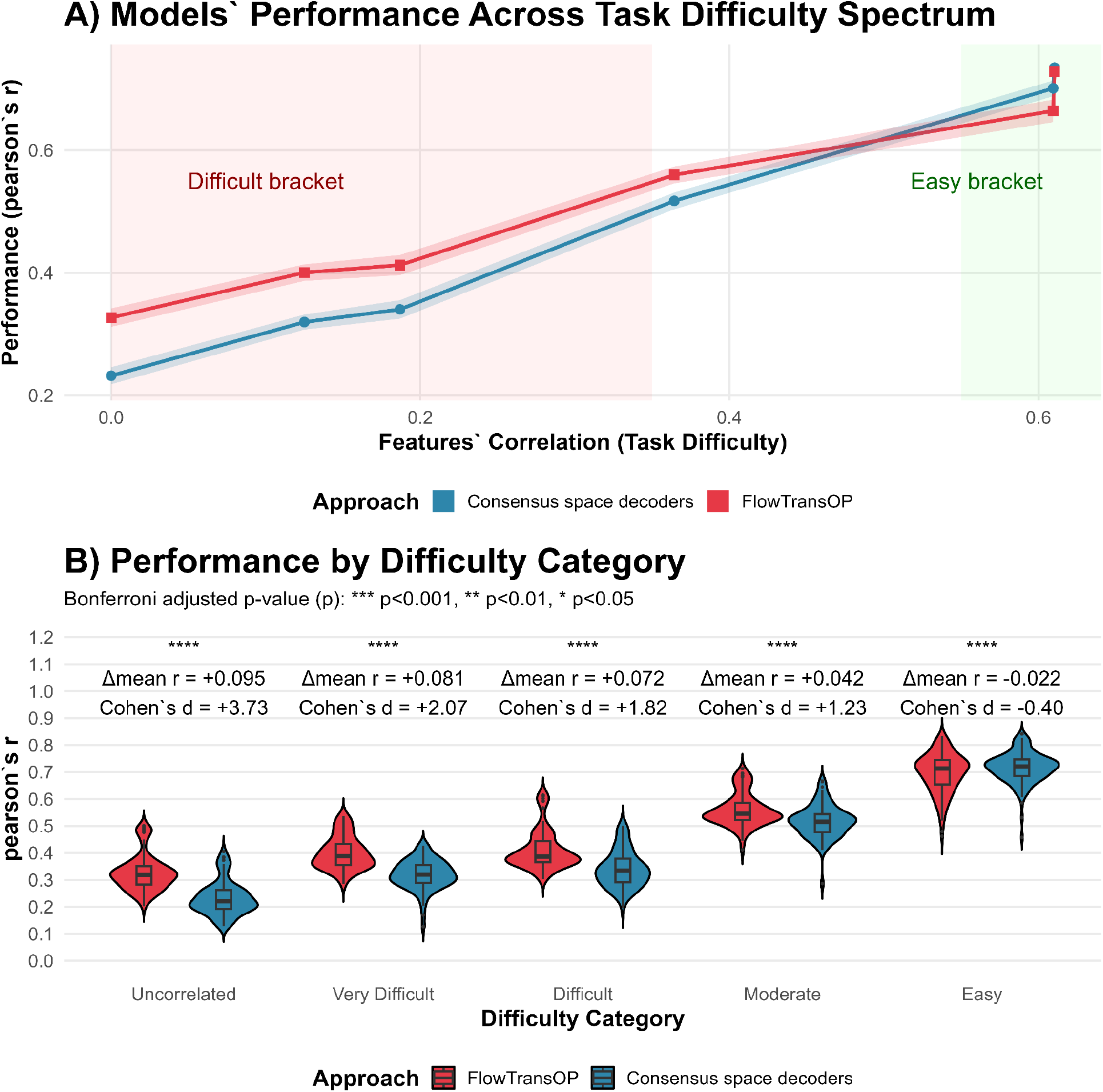
Performance in the hardest regime of highly heterogeneous domains. a) Pearson correlation between predicted and true gene expression values for increasing correlation between input features from the two domains. **b)** Violin plots showing the bracketed performance per difficulty category.

These points highlight the necessity of choosing the correct approach for each difficulty regime of translation. Contrastive learning-based approaches are ideal when there are paired conditions, while in the absence of them, decoders trained on pre-aligned spaces are only good for more homogenous domains (more in section 2.5).

### 2.4. Flow matching is robust to sparse pairing, but decoders suffice when correlations exist

AutoTransOP, as well as other contrastive learning or paired-based approaches, perform poorly as the number of pairs decreases^9,31^. AutoTransOP specifically was shown to lose its predictive power when the amount of paired conditions is less than 10-15% of the training data^9^, which can correspond to even 50-100 paired samples in some cases of large training sets. However, FlowTransOP, which utilizes constrained Flow Matching, is unaffected by the number of paired samples (since it is not using that information) and, in fact, outperforms AutoTransOP in cases with an extremely low number of pairs, in the order of only ~3-7 pairs, while there is no statistical difference in around 34 pairs (Figure 3d). This is the first indication that FM-based approaches can start outperforming contrastive learning in extreme cases, while the first cannot even be applied in the complete absence of pairs. The hybrid approach delivers significantly better performance than the strictly unpaired method when many pairs are available, yet it remains just as robust when pairs are extremely scarce (Figure 3d). Consequently, if researchers are unsure whether they possess enough paired samples, the hybrid strategy represents the optimal balance between the contrastive-based AutoTransOP and the fully unpaired FlowTransOP.

This means that using some approximate pre-alignment and training decoders in the resulting consensus space is the best option for the regime of a low number of pairs but mildly homogenous feature spaces. Even in cases where the input features are different but highly correlated, these approaches are the best option (Figure 4a). However, when looking at hard regimes where no pairs exist, and the features at each domain are different and uncorrelated, FlowTransOP is the only approach that can deliver sufficiently high performance (Figure 4).

### 2.5. Flow matching surpasses baselines in unpaired translation with uncorrelated or distinct features

To fully examine the hardest regimes of highly heterogeneous domains, with no pairs available, we select the 16 cell lines in the L1000 dataset with the most data (for fairness, this is where AutoTransOP’s performance was originally evaluated^9^) and artificially split their gene input space in half randomly 5 times. Each model is evaluated in a 5-fold split, where 80% of the data is used for training and 20% for testing. Since the L1000 landmark genes are generally uncorrelated, randomly splitting results in very uncorrelated features (Supplementary Figure S6), and there, even though FlowTransOP outperforms both the random baseline and the pre-aligned space decoders-only approach (Figure 4, Supplementary Figure S3), its performance is low. This is because this is the hardest possible regime with completely uncorrelated domains and zero pairs. Strikingly, only FlowTransOP achieves meaningful correlations, higher than random baselines (Supplementary Figure S3).

To evaluate most realistically the capabilities of the two unpaired approaches, we use the same data-splitting strategy, but this time making sure we select splits of genes of increasingly high correlation, thus creating easier cases. In the easiest case, where we include some genes in both domains and genes with high correlation, the two approaches perform the same (Figure 4a). However, a clear practical threshold emerges; for domains with feature correlations less than ~ 0.58 (which is still a significant correlation of features), FlowTransOP has the best performance (Figure 4a). This means that for heterogeneous features, even with mild correlation (giving moderate difficulty), FlowTransOP should be selected as the gold-standard approach (Figure 4b).

Only in easy cases with highly correlated features, or sharing the same features between two domains, are the decoders trained on the pre-aligned space outperforming FlowTransOP (Figure 4b). These results demonstrate that FM uniquely enables translation across fully heterogeneous domains where no direct correspondences exist, and provides a potential approach for translating pre-clinical data for novel therapeutics.

### 2.6 FlowTransOP builds a foundational mouse-human transcriptomic map from the ARCHS4 dataset

After benchmarking using the L1000 dataset, we decided to demonstrate the usefulness of FlowTransOP in a real species translation setting, without using any information about pairs. We utilized the ARCHS4 dataset^34^, a transcriptomics dataset containing publicly available RNA-seq data from mouse and human samples. This reflects a fully heterogeneous translational problem, as mouse and human expression matrices have different feature spaces, no paired samples, and only partial biological correspondence through orthologous genes. We therefore trained species-specific autoencoders and bidirectional FlowTransOP velocity fields on ARCHS4 mouse and human bulk transcriptomes, creating a foundational distributional map for translating expression states between species (Figure 5a). Since we do not have ground truth translation labels, the main source of evaluation for this case study is a cyclic approach, where starting from a reference species, we translate gene expression signatures to another target species, and then we translate back to the original reference, to evaluate the consistency of this circular movement between domains (Figure 5b). We especially focus on evaluating held-out liver samples (apart from standard 10-fold cross-validation), as our case study is on Metabolic Dysfunction-Associated Steatohepatitis (MASH).

**Figure 5.**
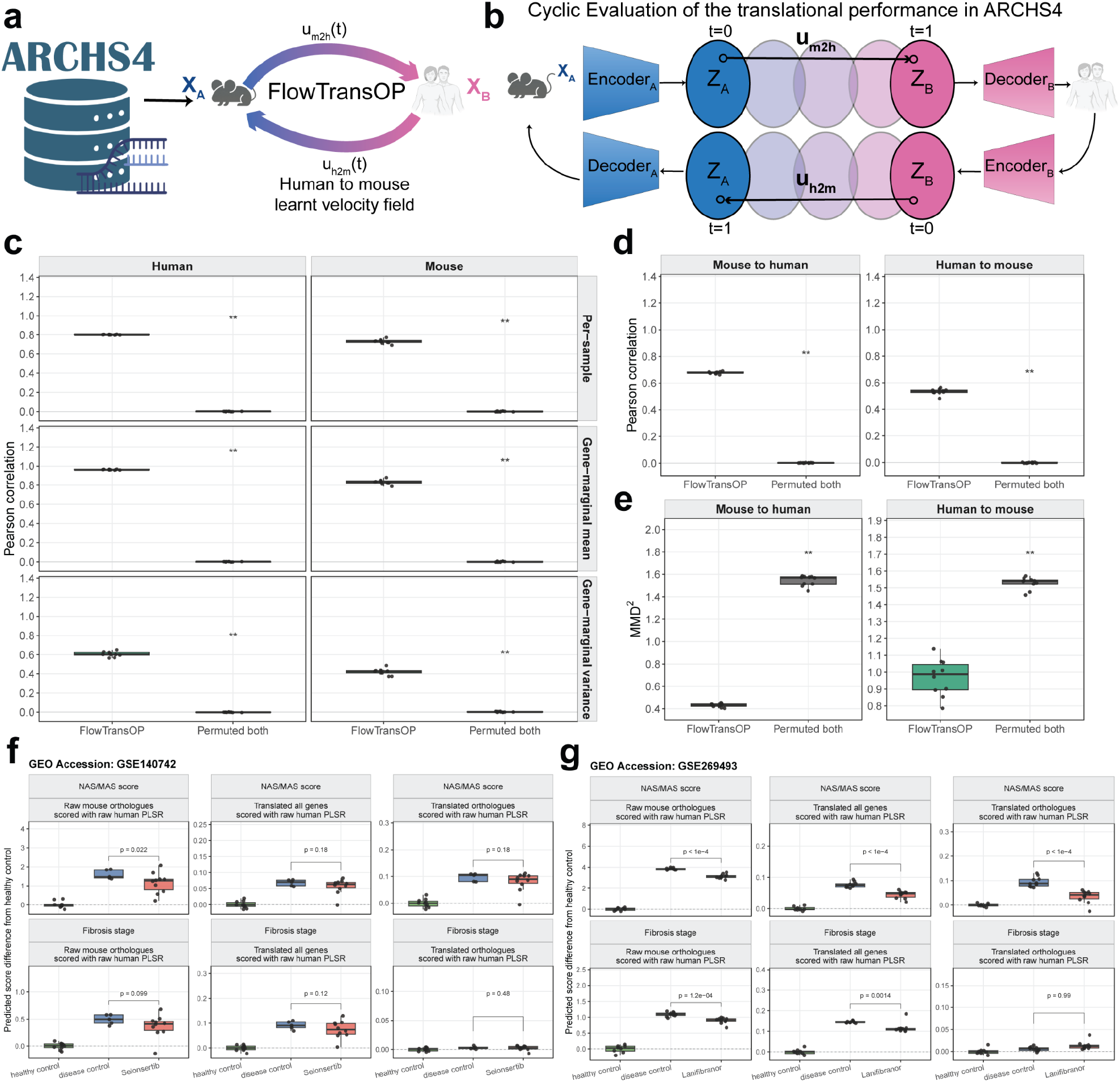
ARCHS4-trained FlowTransOP enables bidirectional mouse-human transcriptomic translation and predicts MASH drug efficacy. **a)** Schematic of FlowTransOP trained on ARCHS4 mouse and human transcriptomes to learn bidirectional velocity fields between species-specific latent spaces. **b**) Cyclic evaluation strategy: Samples are encoded, translated into the opposite species’ latent space, decoded, re-encoded, translated back, and decoded into the original species. **c**) External liver cycle consistency performance across 10 ARCHS4 cross-validation folds. Pearson correlations quantify per-sample reconstruction, gene-marginal mean preservation, and gene-marginal variance preservation after cyclic translation. **d**) Liver orthologue-mediated evaluation. Translated profiles were restricted to matched orthologs and compared with source-side orthologue expression using Pearson correlation. For both c and d, statistical significance was assessed across the 10 folds using paired one-sided Wilcoxon signed-rank tests comparing FlowTransOP against the permuted-both baseline, with the alternative hypothesis that FlowTransOP has a higher Pearson correlation. **e)** Liver latent-space distributional matching, measured by MMD^2^ between translated source liver latents and real target liver latents. Lower MMD^2^ indicates better target-distribution matching. Paired one-sided Wilcoxon signed-rank tests compared FlowTransOP against permuted-both with the alternative hypothesis that FlowTransOP has a lower MMD^2^. **f**,**g**) MASH case-study scoring using PLSR models trained on raw human Govaere expression. Mouse liver perturbation profiles were evaluated as raw mouse orthologues or after FlowTransOP translation into human expression space, and scores are shown as predicted differences from healthy controls. In panel **f)**, results are shown for the GSE140742 dataset (Selonsertib case study). In panel **g**), results are shown from the GSE269493 dataset (Lanifibranor case study). p-values were computed using one-sided Wilcoxon rank-sum tests comparing drug-treated samples against disease controls, with the alternative hypothesis that drug treatment reduces predicted NAS/MAS or fibrosis score. Healthy controls are shown as the zero-reference group for score differences. Asterisks denote adjusted p-values for fold-level comparisons: *p ≤ 0.05, **p ≤ 0.01, ***p ≤ 0.001. In all boxplots, the centerline denotes the median, the bounds of the box denote the 1st and 3rd quantiles, and the whiskers denote points not being further from the median than 1.5 × interquartile range (IQR).

We first evaluated this map by 10-fold cross-validation on held-out ARCHS4 samples. When testing for cycle consistency, FlowTransOP achieved strong recovery for both species, with mean per-sample Pearson correlations of 0.80 in humans and 0.73 in mice. Gene-level marginal structure was also preserved, with mean expression correlations of 0.980 and 0.937 and variance correlations of 0.716 and 0.574 for human and mouse, respectively, while randomized controls (model trained on permuted genes) were near zero (Supplementary Figure S7). We then evaluated orthologue-mediated preservation, investigating the extent to which translated profiles retained the source sample’s matched orthologous expression structure across 15,666 human-mouse orthologs. Even though it is known that orthologs’ expression is not necessarily that similar between the two species, even for the same conditions, we reason that some information do exist, and that when translating with our model, this correlation should be better than random baselines^12,36,37^. FlowTransOP achieved mean per-sample orthologue correlations of 0.557 for human-to-mouse and 0.624 for mouse-to-human translation, again strongly exceeding the permuted baseline (Supplementary Figure S7). Finally, the distributional agreement was evaluated by maximum mean discrepancy, MMD^2^, in both latent and expression space. Lower MMD^2^ indicates that translated source samples better match the target-species distribution. FlowTransOP consistently reduced MMD^2^ relative to the permuted control in both directions (Supplementary Figure S8).

We next tested whether this ARCHS4-trained map generalized to liver samples specifically, an external tissue context relevant to MASH. Before evaluating translation, we confirmed that the species-specific autoencoders retained liver expression information. On held-out liver samples, FlowTransOP autoencoders reconstructed gene-expression means with high performance in both species, with Pearson correlations of 0.975 in human and 0.935 in mouse. Reconstructed gene variances were also correlated with observed variances, with correlations of 0.642 and 0.598, respectively (Supplementary Figure S9). Liver-specific cycle consistency remained strong, with per-sample correlations of 0.801 in human and 0.731 in mouse and gene-marginal mean correlations of 0.960 and 0.832 (Figure 5c). Orthologue-mediated evaluation on liver samples similarly showed preserved cross-species structure, with mean per-sample orthologue correlations of 0.533 for human-to-mouse and 0.677 for mouse-to-human translation (Figure 5d). Latent-space MMD^2^ further showed that translated liver samples were closer to real target-species liver samples than the permuted baseline, both for mouse-to-human translation (0.432 versus 1.544) and human-to-mouse translation (0.968 versus 1.529, Figure 5e). Together, these results indicate that the ARCHS4-trained FlowTransOP map is not only valid in held-out ARCHS4 folds but also preserves biologically meaningful liver structure in an external test setting.

Finally, we used the ARCHS4 mouse-human map for a therapeutic case study in MASH. Mouse liver perturbation signatures were translated into human expression space and scored using Partial Least-Squares Regression (PLSR) models trained on human MASH samples from the Govaere et al. clinical cohort^38^. PLSR has been shown to perform well in this task^39^, but additional Leave-One-Out-Cross-Validation (LOOCV) was performed in this study too (Supplementary Figure S10). Disease scores were reported as predicted differences from healthy controls. In the GSE140742 dataset^40^, mice (Nlrp3A350V mouse strain) were treated with Selonsertib (GS-444217), which is a drug that, even though it was promising in treating liver fibrosis preclinically, did not meet efficacy endpoints in clinical trials^41^. Raw mouse orthologs in the data indeed suggest a modest reduction in NAS/MAS score and fibrosis stage, but after FlowTransOP translation, the predicted human-space effects were not significant for NAS/MAS score or fibrosis stage (Figure 5f). This agrees with the subsequent clinical failure of Selonsertib in phase III NASH/MASH trials, and demonstrates how FlowTransOP could have predicted such failure. In the GSE269493 dataset^42^, samples were obtained from healthy mice and mice under high fat diet (the CDAA-HFD mouse model), as well as CDAA-HFD mice treated with Lanifibranor, an investigational drug associated with MASH with promising results^43^, currently in a Phase 3 clinical trial (NCT04849728)^44^. Lanifibranor retained a stronger predicted therapeutic signal after translation, corroborating its promising results from the literature. Translated all-gene human-space scores predicted significant reductions in both NAS/MAS score (p < 1e-4) and fibrosis stage (p = 0.0014), while translated orthologue scores retained a significant (p < 1e-4) NAS/MAS signal (Figure 5g).

Together, these analyses show that ARCHS4 can be used to train a foundational mouse-human transcriptomic translation map. This map preserves expression structure in cross-validation, generalizes to held-out liver data, and can be coupled to human disease scoring models to make useful clinical predictions.

## 3. Discussion

This study presents FlowTransOP, a generative deep learning framework that uses flow matching techniques to translate between different biological domains without requiring paired training data. We additionally establish guidelines for selecting the optimal translational approach across four difficulty regimes, defined by the degree of feature overlap and sample pairing availability. FlowTransOP aligns the latent distributions of distinct biological systems through distributional matching constrained by a pre-alignment strategy. This enables the translation across heterogeneous contexts without the requirement of similar features, or any direct matching of conditions or samples. FlowTransOP systematically outperforms naive and random baselines while remaining competitive with contrastive methods, or even surpassing them when pairs are scarce, thereby providing a robust, generalized framework for translation across all omic modalities and cases.

The lack of a sufficient number of paired conditions is the main bottleneck of contrastive learning approaches (such as AutoTransOP^9^) in real-world biological applications. While indeed such approaches remain the gold-standard for omics and multi-modal translation^45,46^ (when there are enough pairs), there is a clear advantage of FlowTransOP in scarce pairs settings. However, flow matching techniques by themselves do not guarantee that the transformation between domains is done effectively, i.e., preserving the meaningful structure of the data. Thus, initial pre-alignment to act as a guide for the transformation is necessary, and as indicated in this study, the quality of the initial pre-alignment is crucial. Moreover, the pre-aligned space itself is informative enough to provide high-fidelity omics translation between biological domains (even higher than FlowTransOP), but performance drops dramatically when the input feature spaces are more dissimilar. FlowTransOP outperforms the pre-aligned space even in cases of moderate similarity between the input domains. No single method, however, is universally the best, highlighting the need to identify into which translational regime each case study falls. Nevertheless, as demonstrated in this study, in cases of uncertainty regarding the similarity of the input features and the amount of paired conditions, FlowTransOP, utilizing a hybrid constraining approach (using both the pre-aligned space and available pairs), should be chosen as the universally best intermediate solution.

FlowTransOP effectively decodes the underlying structural consistency of heterogeneous biological distributions, providing a robust and reliable engine for therapeutic inference when traditional orthology-based and contrastive methods fail. Realistically, in novel therapeutics’ development, where there are not yet paired conditions between humans and pre-clinical models, researchers can utilize general data from different sources for each domain to build a flow from pre-clinical model to humans, enabling them to estimate the potential effect of a therapeutic modality on patients, in terms of a selected molecular signature, before an actual clinical trial. This allows for early adjustments, reducing the time and cost of therapeutic development, especially for new emerging diseases (e.g., COVID-19 pandemic^47^). The ARCHS4 case study illustrates this intended use case in a realistic cross-species setting. By training on large unpaired mouse and human transcriptomic collections, FlowTransOP learned a foundational mouse-human expression map that generalized to held-out liver data and enabled mouse perturbation signatures to be interpreted in human MASH-associated score space. The Selonsertib and Lanifibranor examples are not intended as definitive drug-development claims, but they show how human-centered translation of preclinical perturbation data could help prioritize therapies before costly clinical testing. This aligns with a broader emerging view that preclinical perturbation experiments are most useful when they can be read in human biological coordinates rather than only in the native coordinate system of the model organism^16^.

Despite its promise as a generalized translation model, establishing that it is possible to translate between biological systems using generative deep learning with virtually zero requirements, FlowTransOP has its own limitations. First of all, performance declines in the hardest regimes with uncorrelated features and no pairs, and results depend heavily on the quality of initial pre-alignment. Even though the performance is always higher than random and naive baselines, it is still practically low in the hardest possible case. This means that FlowTransOP can be used as the baseline standard for comparing the performance of other models, as researchers work on building more reliable translation models. Additionally, performance is constrained by our ability to train a high-performing autoencoder for each domain. In cases of low data availability, with simultaneously very high dimensionality (in the order of thousands of molecular features), this may prove to be a challenge. Fortunately, pretraining on larger datasets (such as ARCHS4^34^ for humans and mice, and Tahoe-100M^48^ for single-cell data) and fine-tuning approaches can alleviate this problem. A natural extension involves the deployment of recent large foundational biological models^49–54^, instead of training separate autoencoders for each biological system, or creating new such models using larger datasets.

All these demonstrate that recent advances in deep learning have now enabled the translation between biological systems, fully unconstrained. In this study, we focused on aligning the starting (source) and ending (target) distributions. In this first implementation, we intentionally used a simple conditional optimal-transport, or linear, probability path with a fixed-step midpoint ODE solver. This choice makes the model stable and interpretable, but it does not exhaust the FM design space. Thus, a compelling future direction is the explicit optimization of the actual transport path of the ODE itself, exploring richer probability paths, alternative couplings, stochastic bridges, or biologically constrained trajectories. In other fields, such as robotics, trajectory-optimized density control is used to enforce critical path-dependent properties^55^. FlowTransOP could optimize the ODE trajectory to enforce biological constraints along the intermediate steps of the transport path, or even learn to build trajectories for longitudinal omic profiles. This would ensure that as the model continuously translates a signature from mice to humans, the intermediate latent states represent biologically viable or plausible intermediate omics profiles, rather than just abstract mathematical transitions. In combination with recent advances in Large Language Models (LLMs), which can be utilized to explore the prior knowledge space and interpret the translated outcome, FlowTransOP and similar approaches are sure to make pre-clinical therapeutic and immunization design more efficient and effective.

## 4. Materials and Methods

### 4.1. Data acquisition and preprocessing

To ensure fairness when comparing with the state-of-the-art, the initial, pre-processed L1000^35^gene expression data were accessed and downloaded from the GitHub repository of AutoTransOP^9^: https://github.com/Lauffenburger-Lab/OmicTranslationBenchmark. Additionally, the performance of AutoTransOP in translating the 978 landmark genes’ expression profiles of the L1000, for some tasks, was also retrieved from the same repository. The tasks’ performance that was loaded for the gold-standard were: i) the performance in translation for different sample sizes for 8 cell line pairs, and ii) the performance in translating artificially different biological domains, where the landmark genes are split in half and used with two different autoencoders for 16 different cell lines. For a fair comparison, FlowTransOP was trained on the same splits. For the process followed to extend our benchmark to domains with different similarity between their input features (the one loaded from the GitHub repository is completely dissimilar and distinct features) see more details in section 4.2. For the process followed for cases with extremely few pairs see section 4.3.

### 4.2. Extension of splits on artificially heterogeneous domains of different similarity levels

We generated more subsets of our features to expand our analysis on different brackets of similarity of features between input domains. Specifically, we generated gene subsets of increasingly high inter-domain feature similarity, creating a bracketed spectrum of domain heterogeneity. First, the retrieved, full L1000 gene expression matrix (including measured landmark genes and computationally inferred genes of the L1000 dataset) was used to compute a gene-by-gene Pearson correlation matrix across all available samples. The top 978 genes by mean inter-gene correlation were selected as the working gene space, and a softmax-weighted probability distribution over their mean correlation scores was derived to bias random sampling towards more mutually correlated genes. An iterative random search (with a maximum of 1000 iterations) was then performed to identify the partition of these 978 genes into two equally-sized subsets that maximised the mean cross-correlation between the two halves, constituting the “best” bracket. To encourage broader exploration of the solution space, the softmax weights were relaxed to uniform after 200 iterations whenever the relative improvement in mean cross-correlation dropped below 5%. This bracketing procedure was repeated sequentially on the remaining genes (excluding those already assigned to a bracket) to obtain the second, third, and fourth best brackets in descending order of inter-domain feature similarity, yielding four difficulty levels overall. To extend the spectrum towards the highest feature similarity regime, an additional class of gene subsets was constructed by introducing deliberate feature overlap between the two domains. Specifically, starting from the best bracket, 30% of the genes in each subset were randomly exchanged with genes drawn from the complementary subset, such that a controlled fraction of features became literally shared between domains. This procedure was repeated for 5 random iterations, producing 5 high-correlation gene pairs representing the easiest translational regime. The samples used in the 5-fold CV were the same ones that were retrieved from the GitHub repository.

### 4.3. Benchmark on an extremely low number of paired samples

To further investigate performance in the most data-scarce regime, we evaluated all models on an extremely reduced number of paired training samples, using the A375–HT29 cell line pair (for fairness with what was used in AutoTransOP^9^). Specifically, experiments were conducted with only 1, 2, or 3 paired samples available during training, while the full unpaired data for each domain were retained. For each paired-sample count, up to 30 non-overlapping repeats were drawn per cross-validation fold by randomly permuting all available training pairs and partitioning them into consecutive disjoint subsets of the target size, ensuring that no two repeats within the same fold shared a paired sample.

### 4.4. Autoencoder architecture and pre-training for each domain

For each biological domain (i.e., cell line, or potentially different species), we train its own separate autoencoder, so that we can embed the input gene expression (or any omic profile) in latent spaces of the same dimensionality. Later translation via flow matching is performed between these dimensionally reduced spaces. Autoencoders are pre-trained to minimize the mean squared error in reconstruction (summed across features) for 1000 epochs, with a batch size of 120 (except for the artificially different domains benchmark, where a batch size of 512 is used), using the Adam optimizer with a learning rate of 0.001. Additionally, the L2-regularization is used to penalize large weights (λ=0.01). Both the encoder and decoder use the same architecture as AutoTransOP’s autoencoders, which consist of 3 sequential components, including a fully connected layer, followed by a batch normalization layer (with 0.6 momentum), an ELU activation function, and a dropout layer (except for the final component, which has only a fully connected layer). The encoder reduces the dimensions to 640, 384, and ultimately a latent dimension of 292, and then the decoder increases the dimension in the reverse order. Only in the case of different input domains, the dimensions change to 384, 256, and 128, respectively, as the input dimension is half. The decoder utilizes a dropout rate of 0.2, and the encoder a dropout rate of 0.1, with an additional dropout layer with a rate of 0.5 for the input profile directly. Finally, a scheduler reduces the learning rate by 20% every 200 epochs. All these are similar hyperparameters to the ones used in AutoTransOP, for case studies with similar dimensionality of the data.

### 4.5. Pre-alignment of domains

To pre-align the two input domains, we adapt the TRANSACT method^23^. Specifically, we developed both a GPU and CPU implementation of TRANSACT (and tested that for many different distributions, CPU and GPU implementations give virtually the same result in Supplementary Figures S11-S19). Briefly, TRANSACT projects both domains into a consensus common space, by separately standardizing each domain, computing centered within- and cross-domain RBF kernels, extracting domain-specific kernel principal components, and aligning them through SVD-derived principal vectors. For each aligned component pair, source and target sample scores were gradually interpolated between the matched source and target directions by intensifying the interpolation point that minimizes the Kolmogorov-Smirnov distance between domains.

Using this pre-aligned space, we calculate the pair sample Pearson’s correlation in the consensus space. This will be later used to construct a mask for constraining the flow matching, so that samples very similar in the pre-aligned space are slightly closer in the post-transferred space, after transferring from one domain to another via flow matching (details in section 4.7).

For the baseline model that just utilizes this pre-aligned space, decoders are trained to take as input the projected samples in the input space and reconstruct the original feature space (gene expression). The same hyperparameters are used as the ones for the decoders during autoencoder training, except for the latent dimension, which uses the dimensions of the consensus space (which, following the TRANSACT original publication^23^, is set to 30).

### 4.6. Flow matching generative model (FlowTransOP)

For transforming a source distribution (e.g., cell line A, or species A, etc.) to a target distribution (e.g., cell line B, or species B, etc.), FlowTransOP (Figure 2) utilizes Flow Matching (FM) to learn a velocity field (*u*_*t*_) to enable the transformation from a source distribution to a target distribution^32^. In FlowTransOP, we parameterize the velocity field using a neural network (*u*_*θ*_). Assuming it has learnt the ground truth velocity field (see section 4.7 for training), it represents the instantaneous speed and direction at every point, and thus it also defines the flow (*ψ*_*t*_) which maps each sample from its starting position to its ending position t (e.g., if *z*_1_~*p*_1_ *then ψ*_2_(*z*_2_)~*p*_2_, where *p*_1_ and *p*_2_ are the source and target/end distributions) are in Eq. 1:

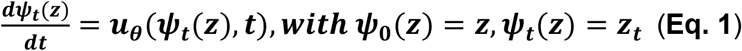

Solving this ODE gives the flow for a transition of interest, while the flow generates the probability path that transforms a source distribution (e.g., cell line A) to a target distribution (e.g., cell line B). For a specific pair of latent representations (*z*_1_,*z*_2_) from different domains, in FlowTransOP, we use the conditional optimal-transport flow to define the transition between domains in Eq. 2:

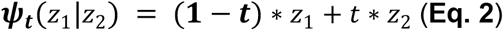

This process defines the flow in the simplest possible way (a straight line between *z*_1_ and *z*_2_), where at t= 0 we are at the source distribution and at the final step (t=1) we are the target distribution. Everything in between is a linear interpolation. When using enough intermediate transitions to move from source to target (here, for speed, we use 10 steps), this definition is accurate enough, though we expect more complex flows may improve translation between domains.

Combining Eq.1 and Eq. 2 results in defining the ground truth velocity field in Eq. 3:

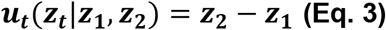

This will be used during training to optimize the neural net parameterizing the velocity field (*u*_*θ*_). During inference, we can integrate over ***u***_*θ*_(***z***_***t***_, ***t***) to move from source (*z*_1_) to target (*z*_2_) distribution with: 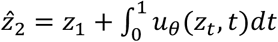. In practice, we approximate this using a second-order midpoint ODE solver, where:

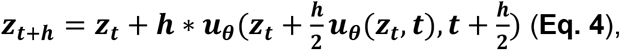

over N uniform steps from t=0 to t=1, and with h=1/N.

### 4.7. Training unpaired flow matching

During training, there are two major parts of the training loss: **i)** the regression objective to learn the velocity field (*L*_*CFM*_), and **ii)** the structural regularization via pre-aligned similarity (*L*_*struct*_). The first loss is minimized through the mean square error minimization of the ground truth velocity vector (Eq. 3) and the parametrized velocity vector *u*_*θ*_(*z*_*t*_, *t*) via a neural network:

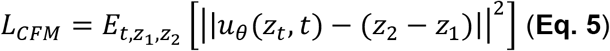

To maintain some structure during flow matching, we pre-align (see section 4.5) samples from the two domains in each batch of training data, and we obtain a max-normalized correlation matrix (*C*) describing the similarity of samples in the batch. Then we use the inference step of FlowTransOP (section 4.6) to transform the source latent representation (*z*_1_) to the target representations 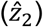, and calculate euclidean distance between samples in the batch (in the target domain only). We aim to softly push samples after transformation to be close to each other if they were very similar in the pre-aligned space (thus, we do not care about negative values of the similarity matrix), so we define the structural loss in the Eq. 6:

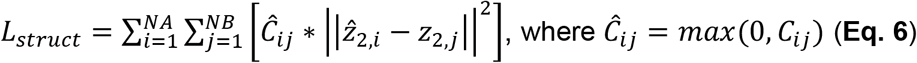

The final total loss during training is calculated as *L* = *λ*_*CFM*_ * *L*_*CFM*_ + *λ*_*struct*_ * *L*_*struct*_, with *λ*_*CFM*_ = 1 and *λ*_*struct*_ = 10^−3^.

### 4.8. Hybrid approaches

For hybrid approaches of FlowTransOP that utilize both available pairs and the pre-alignment for unpaired translation, we redefine *Ĉ*_*ij*_ to guide the transition, by combining the pre-aligned correlation matrix and the known pairs in three variations: **i)** we use the sum (Eq. 7) of the values of the paired binary matrix (denoted as A) (it has 1 if two samples are paired) and the previous correlation matrix (only each positive values), **ii)** we use the mean of those matrices (Eq. 8), and **iii)** we use the max between binary and correlation matrix (Eq. 9).

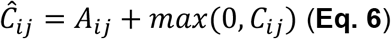

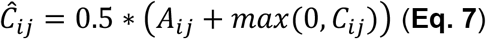

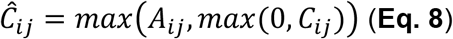

This newly defined similarity matrix Ĉ_*ij*_ is what is now used in *L*_*struct*_ in Eq. 6.

## Data availability

The study did not produce any new experimental data. The preprocessed L1000 data can be accessed and downloaded from the GitHub repository of AutoTransOP^9^: https://github.com/Lauffenburger-Lab/OmicTranslationBenchmark. All analyzed data that were used to train our models and produce all tables and figures are available at https://github.com/NickMeim/FlowTransOP.

## Code availability

The code to generate and pre-process data, figures, and tables, as well as train models, is available in a detailed GitHub repository: https://github.com/NickMeim/FlowTransOP. We organized the FlowTransOP implementation as a package-like repository with modules for preprocessing, model training, translation, benchmarking, and plotting. Users can train FlowTransOP models on their own domains with custom hyperparameters, reproduce the benchmark datasets used here, or apply the same evaluation framework, including reconstruction, cycle consistency, orthologue preservation, MMD^2^, and downstream disease-scoring analyses, to their own models. All instructions are present in the GitHub repository.

## Acknowledgements

This work was funded from NIH NIAID contract #75N93019C00071 and grant AI-181898, as well as the Army Institute for Collaborative Biotechnologies collaborative agreement W911NF-19-2-0026. NM would also like to acknowledge funding from the 2024-2025 Takeda Fellowship.

