## Supplementary Material for "FlowTransOP: Distributional Translation of Omics Signatures via Constrained Deep Flow Matching"


#### Supplementary Figures

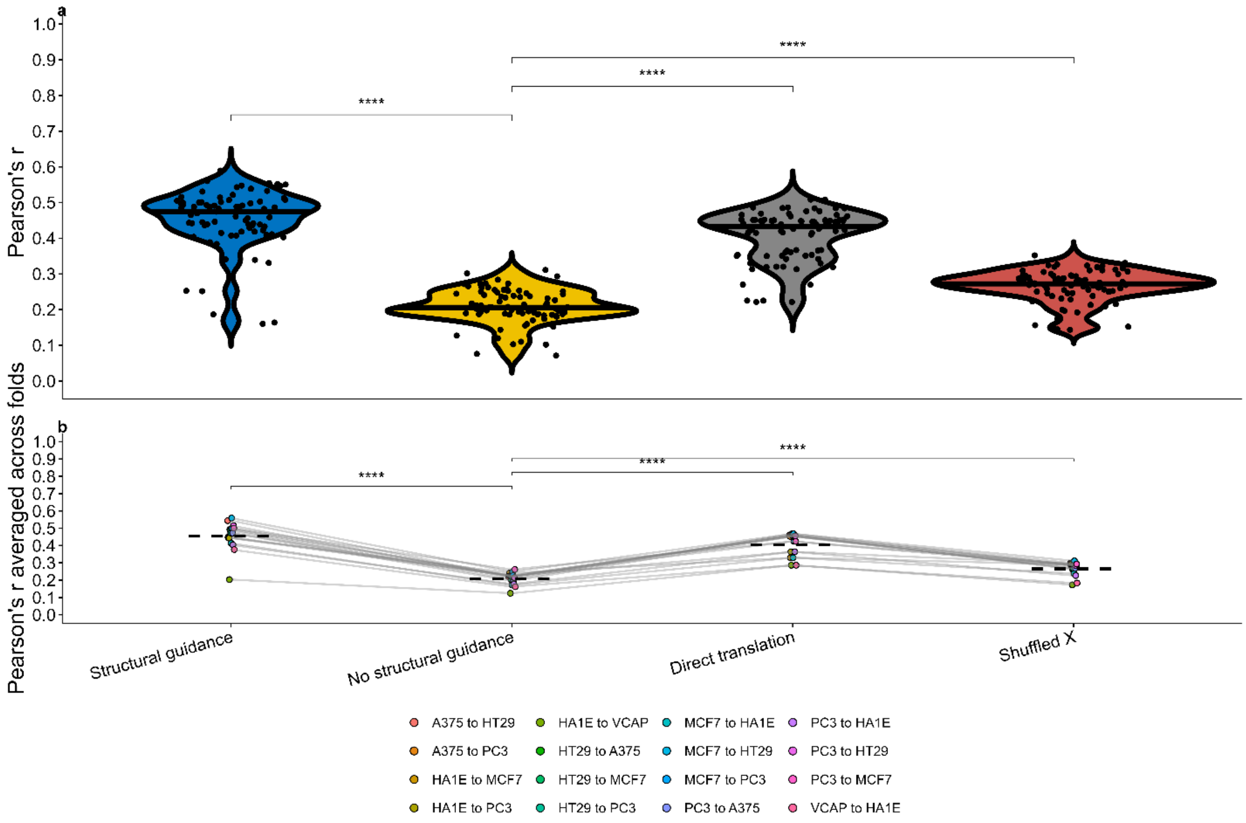


**<u>Supplementary Figure S1:</u> Structural guidance is required for effective FlowTransOP translation. a)** Fold-level translation performance. Violin plots show Pearson correlation between predicted and observed expression profiles for FlowTransOP with structural guidance, FlowTransOP without the TRANSACT-derived structural guidance term, direct translation, and a shuffled features (shuffled X) random baseline.

Points denote individual cross-validation fold and directed cell-line-pair values; the horizontal line within each violin denotes the median. **b)** Same comparison after averaging Pearson correlation across folds for each directed cell-line pair. Each dot denotes one directed cell-line pair, colored by pair, and gray lines connect the same pair across approaches. Dashed crossbars denote the mean across directed cell-line pairs. Only translation directions present in the no-structural-guidance run were included in the matched comparison. p-values were computed using paired two-sided Wilcoxon signed-rank tests: in panel a across matched fold and translation-direction values, and in panel b across matched fold-averaged translation directions. Asterisks denote p-values: *p ≤ 0.05, **p ≤ 0.01, ***p ≤ 0.001, ****p ≤ 0.0001.

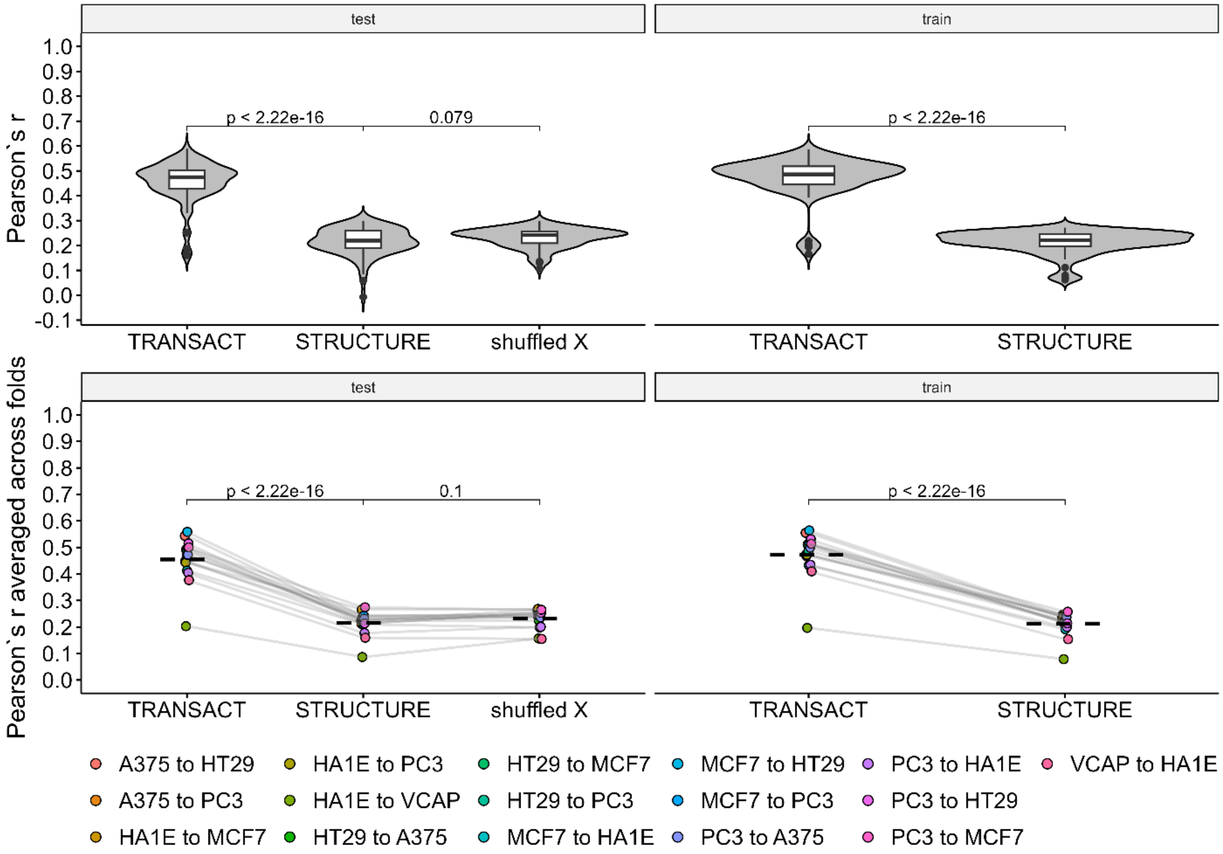


**<u>Supplementary Figure S2</u>: TRANSACT-derived pre-alignment provides stronger structural guidance for L1000 FlowTransOP than STRUCTURE-derived pre-alignment.** Top panels show fold-level Pearson correlation (r) between predicted and observed L1000 perturbational expression profiles for held-out test and training folds after constraining FlowTransOP with either TRANSACT or STRUCTURE; a shuffled-feature control is included for the test set. Bottom panels show the same comparison after averaging r across folds for each directed cell-line translation task, with lines connecting the same task across approaches. Displayed p-values were computed using two-sided Wilcoxon tests comparing the indicated approaches. In all violin/boxplots, the centerline denotes the median, the box denotes the 1st and 3rd quartiles, and whiskers denote points within 1.5 x interquartile range (IQR).

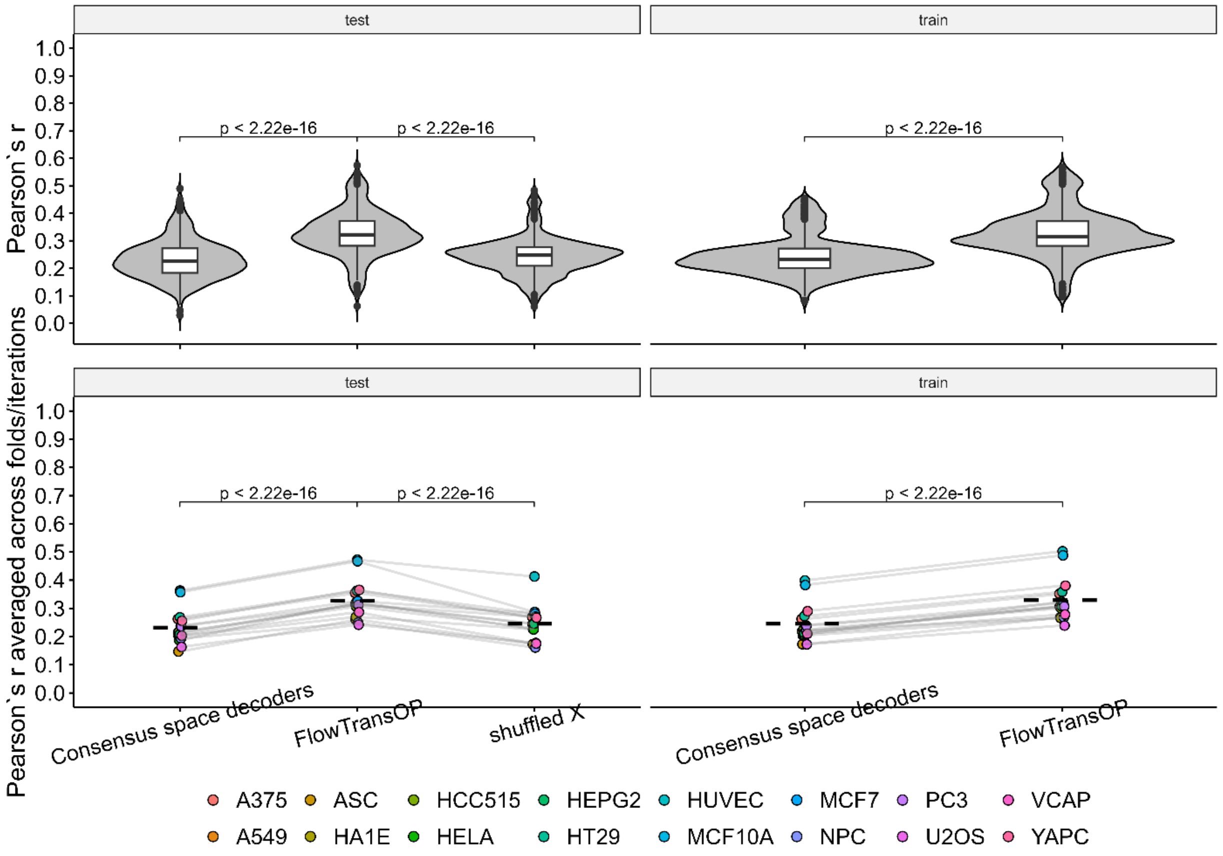


**<u>Supplementary Figure S3:</u> FlowTransOP retains predictive signal in heterogeneous L1000 translation tasks with distinct feature spaces.** Consensus-space decoders, FlowTransOP, and shuffled-feature controls were compared after splitting L1000 genes into non-overlapping input spaces. Top panels show Pearson correlation (r) distributions across folds and iterations for training and held-out test data. Bottom panels show cell-line-level averages, with points connected across methods for the same cell line. p-values shown in the panels were computed using two-sided Wilcoxon tests comparing the indicated approaches. In all violin/boxplots, the centerline denotes the median, the box denotes the 1st and 3rd quartiles, and whiskers denote points within 1.5 x IQR**.**

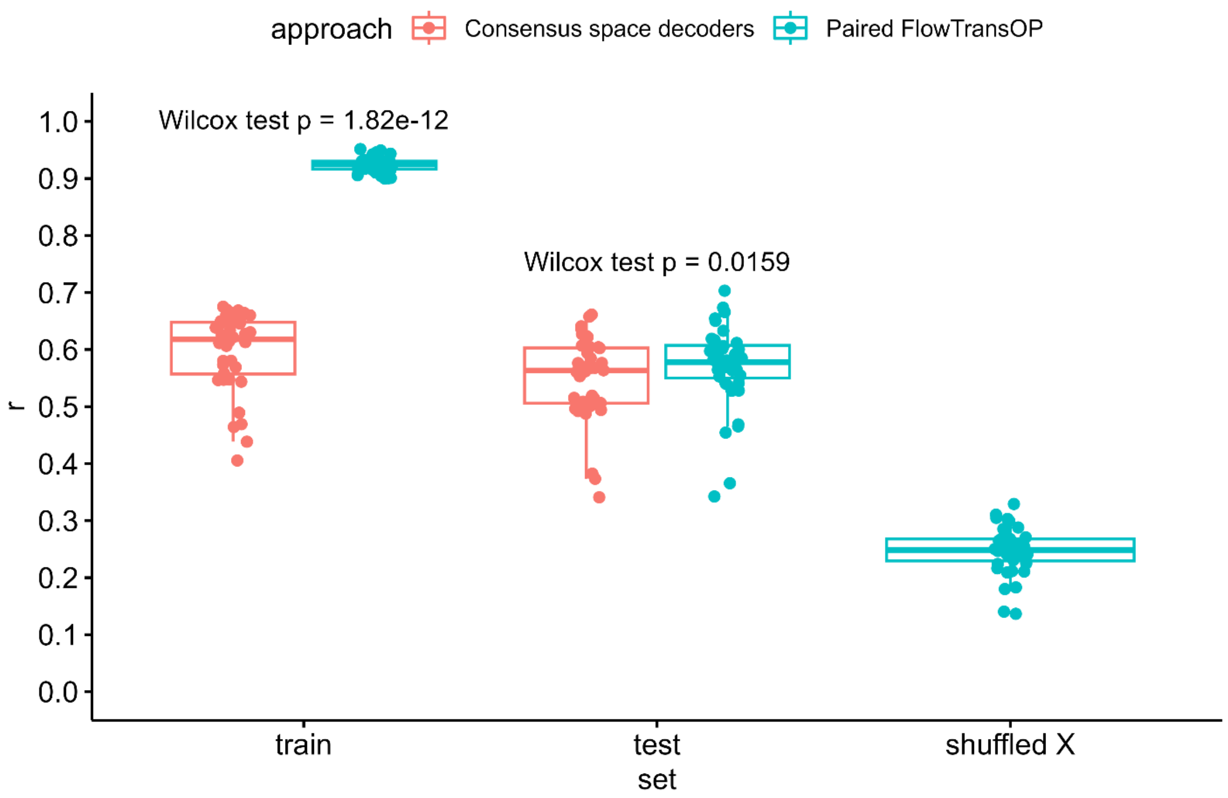


<u>Supplementary Figure S4:</u> **Paired FlowTransOP fits paired training data strongly but provides limited held-out improvement over consensus-space decoders.** Pearson correlation (r) is shown for training folds, held-out test folds, and shuffled-feature controls in the shared-feature paired L1000 benchmark. p-values shown above the train and test comparisons were computed using paired two-sided Wilcoxon signed-rank tests comparing paired FlowTransOP against consensus-space decoders across matched folds/tasks. In all boxplots, the centerline denotes the median, the box denotes the 1st and 3rd quartiles, and whiskers denote points within 1.5 x IQR**.**

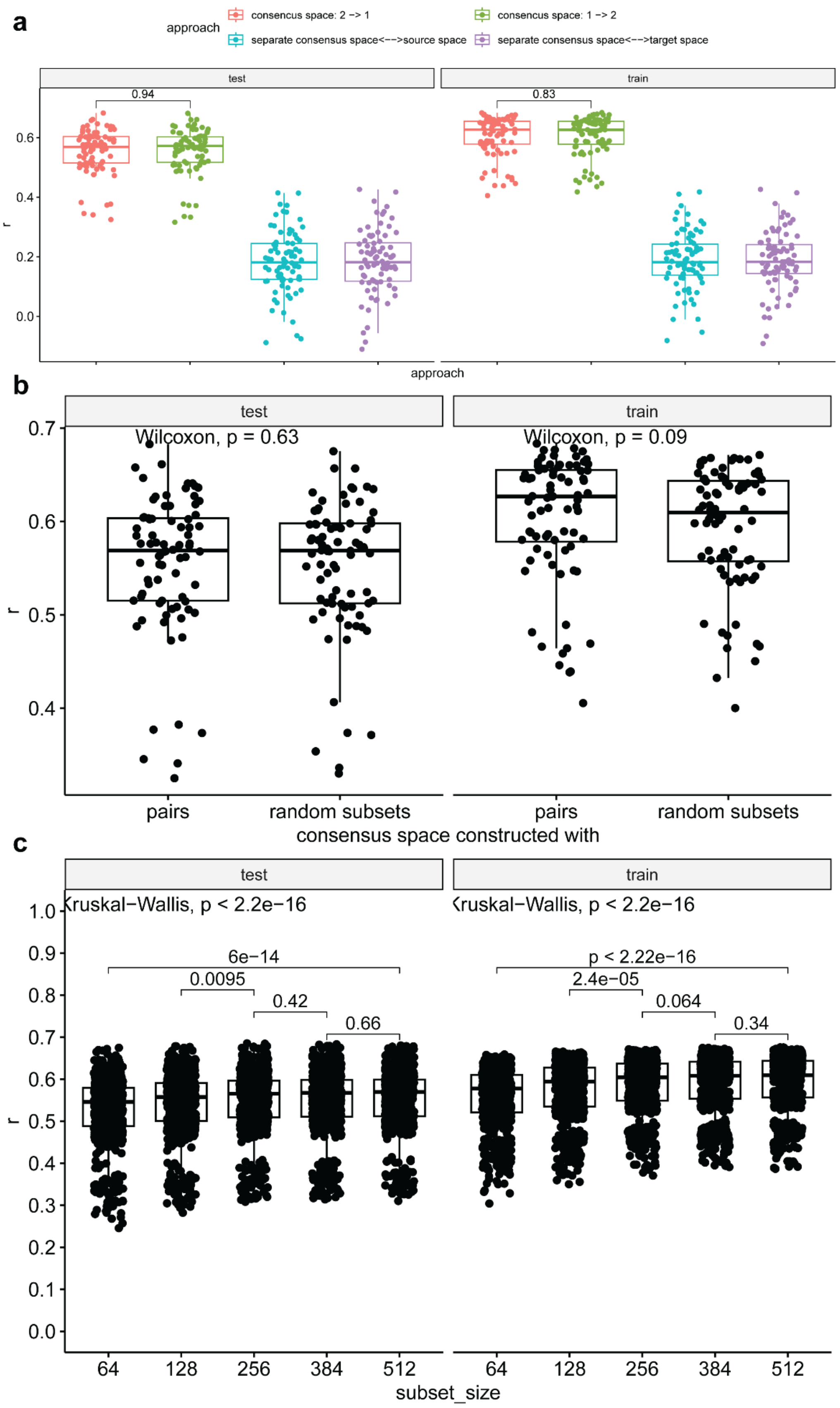


**<u>Supplementary Figure S5:</u> Construction of the pre-aligned consensus space affects downstream decoder performance. a)** Decoder performance when the TRANSACT consensus space was constructed using both source-target orientations or using separate source-only and target-only consensus spaces. Pearson correlation (r) is shown for train and held-out test folds; p-values compare the two shared-consensus orientations using paired Wilcoxon signed-rank tests**. b)** Decoder performance when the consensus space was constructed from known pairs or from random unpaired subsets. p-values were computed using two-sided Wilcoxon tests comparing paired and random-subset consensus construction within each split. **c)** Effect of random-subset size on decoder performance. Global p-values were computed using Kruskal-Wallis tests, and displayed pairwise p-values were computed using two-sided Wilcoxon tests. In all boxplots, the centerline denotes the median, the box denotes the 1st and 3rd quartiles, and whiskers denote points within 1.5 x IQR.

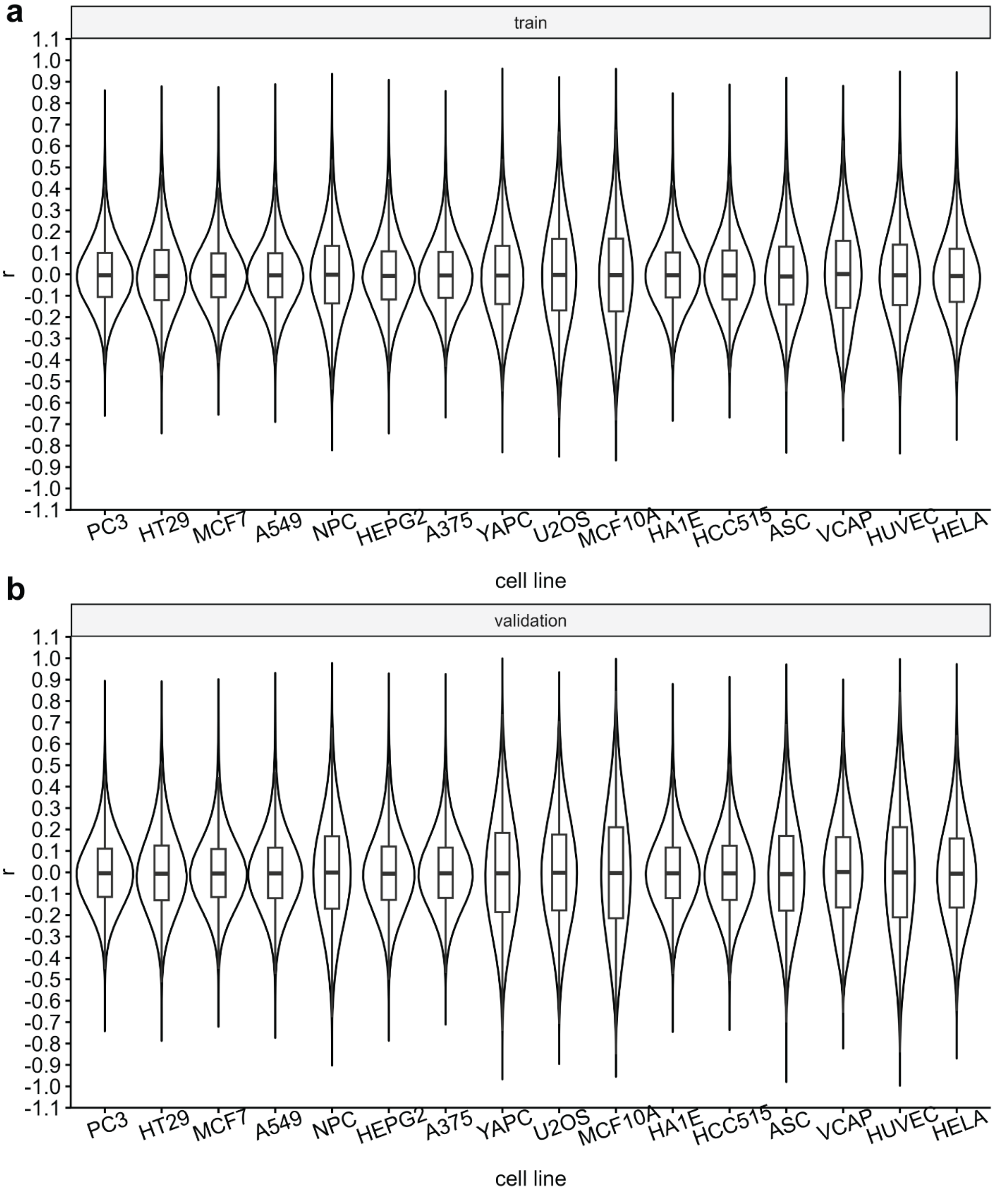


**Supplementary Figure S6: Artificially split L1000 feature spaces span a broad range of inter-domain feature similarity. a)** Distribution of feature-feature Pearson correlations (r) in training splits for each cell line. **b)** Distribution of feature-feature Pearson correlations (r) in held-out validation splits for each cell line. Violin plots summarize the full correlation distribution, with embedded boxplots showing the median and interquartile range. These feature-correlation distributions define the heterogeneous-input difficulty axis used to benchmark FlowTransOP against consensus-space decoders.

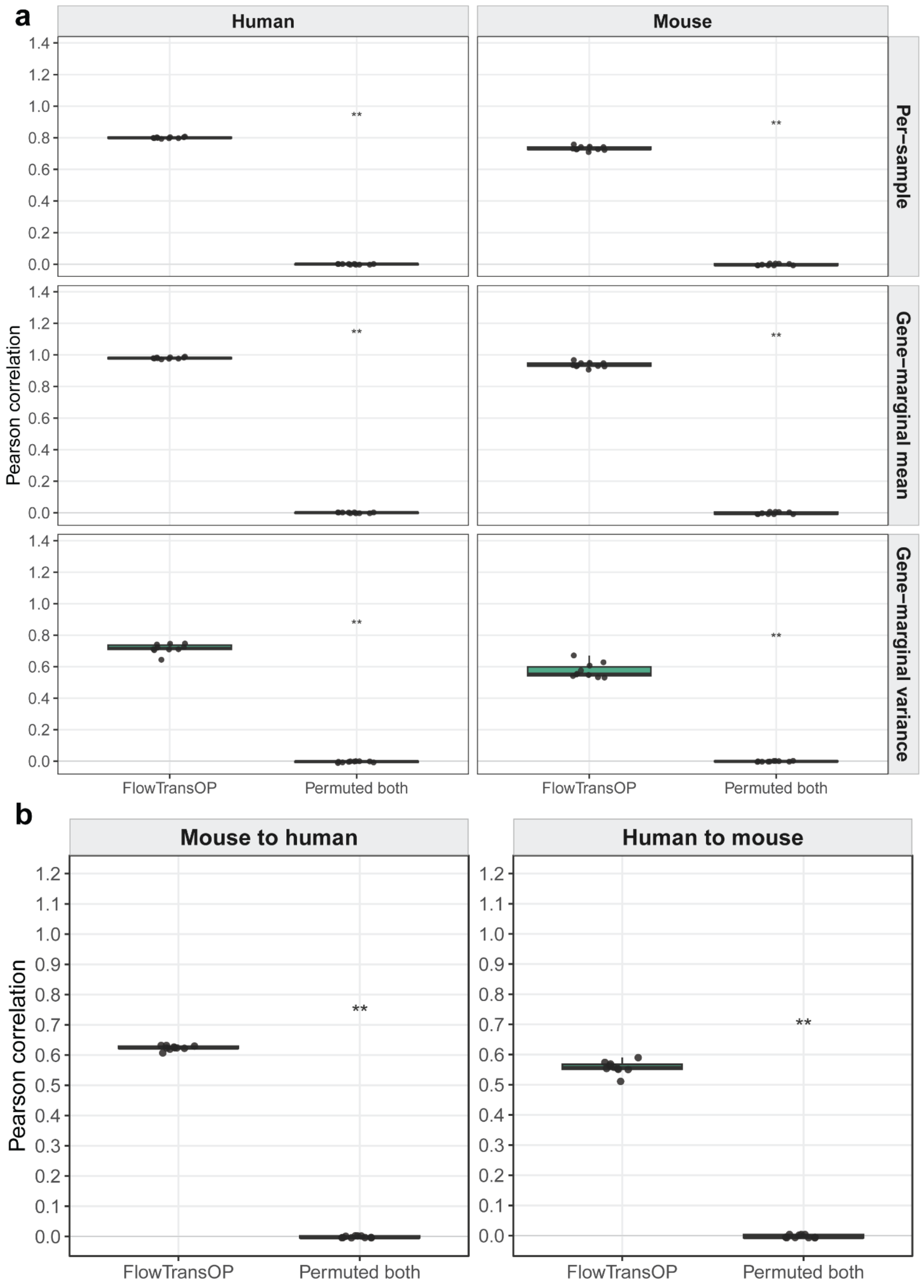


<u>Supplementary Figure S7:</u> **ARCHS4 10-fold cross-validation demonstrates cycle consistency and orthologue preservation. a)** Cycle consistency across held-out ARCHS4 folds. Samples were translated

from the source species into the opposite species and back, then compared with the original source profiles using Pearson correlation for per-sample expression, gene-marginal means, and gene-marginal variances. **b)** Orthologue-mediated evaluation across held-out folds. Translated profiles were restricted to matched human-mouse orthologues and compared with source-side orthologue expression using Pearson correlation. For a and b, statistical significance was assessed across folds using paired one-sided Wilcoxon signed-rank tests comparing FlowTransOP with the permuted-both baseline, with the alternative hypothesis that FlowTransOP has higher Pearson correlation. Asterisks denote Holm-adjusted p-values: *p <= 0.05, **p <= 0.01, ***p <= 0.001. In all boxplots, the centerline denotes the median, the box denotes the 1st and 3rd quartiles, and whiskers denote points within 1.5 x IQR.

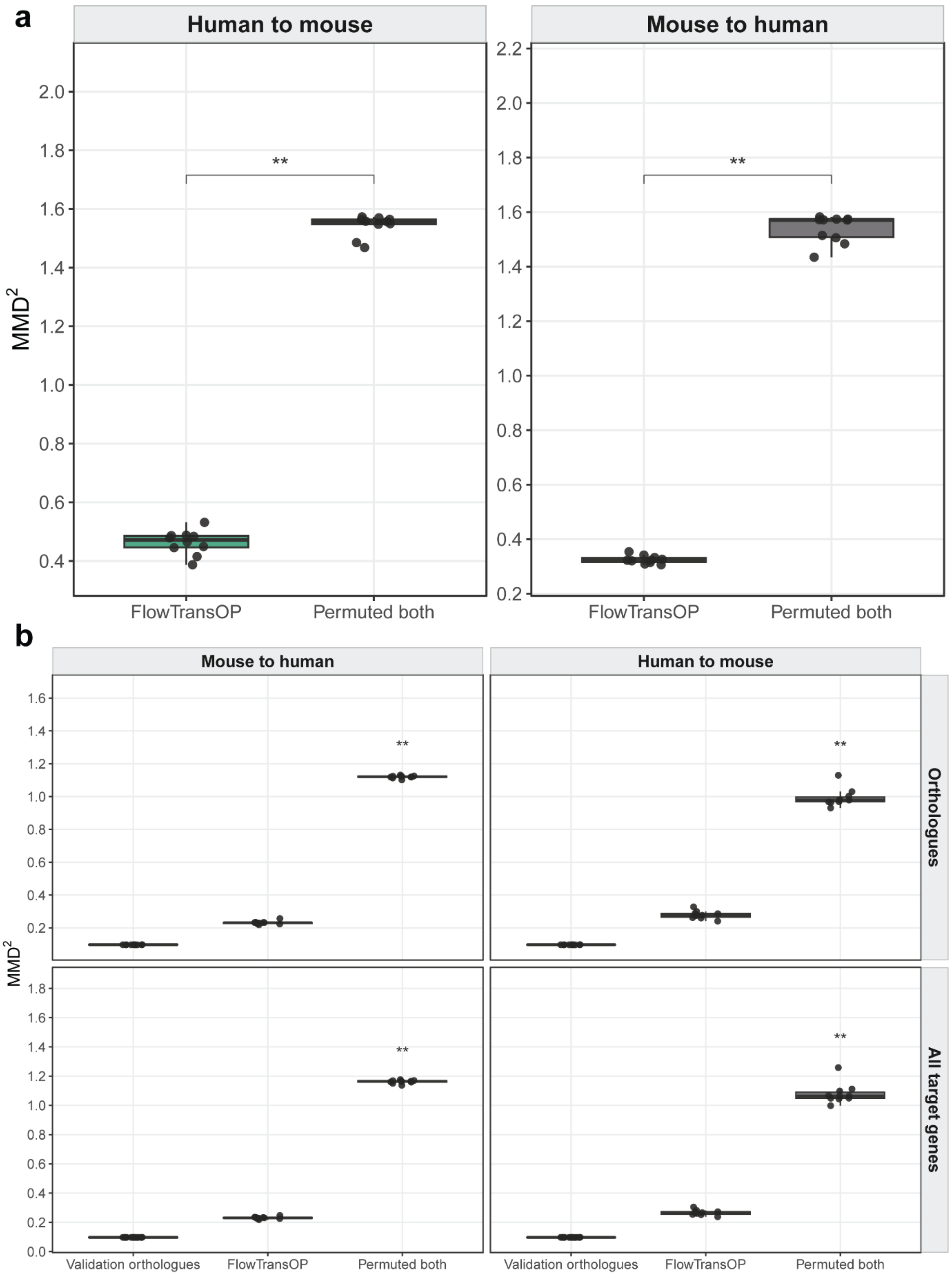


**Supplementary Figure S8: ARCHS4 10-fold cross-validation shows improved distributional matching in latent and expression space. a)** Latent-space distributional matching measured by $MMD^2$ between translated source samples and real target-species samples. **b)** Expression-space distributional matching measured by $MMD^2$ after decoding translated samples, with evaluations over matched orthologues and all target genes. Lower $MMD^2$ indicates better agreement with the target-species distribution. Paired one-sided Wilcoxon signed-rank tests compared FlowTransOP with the permuted-both baseline across folds, with the alternative hypothesis that FlowTransOP has lower $MMD^2$. Asterisks denote Holm-adjusted p-values: *p <= 0.05, **p <= 0.01, ***p <= 0.001. In all boxplots, the centerline denotes the median, the box denotes the 1st and 3rd quartiles, and whiskers denote points within 1.5 x IQR.

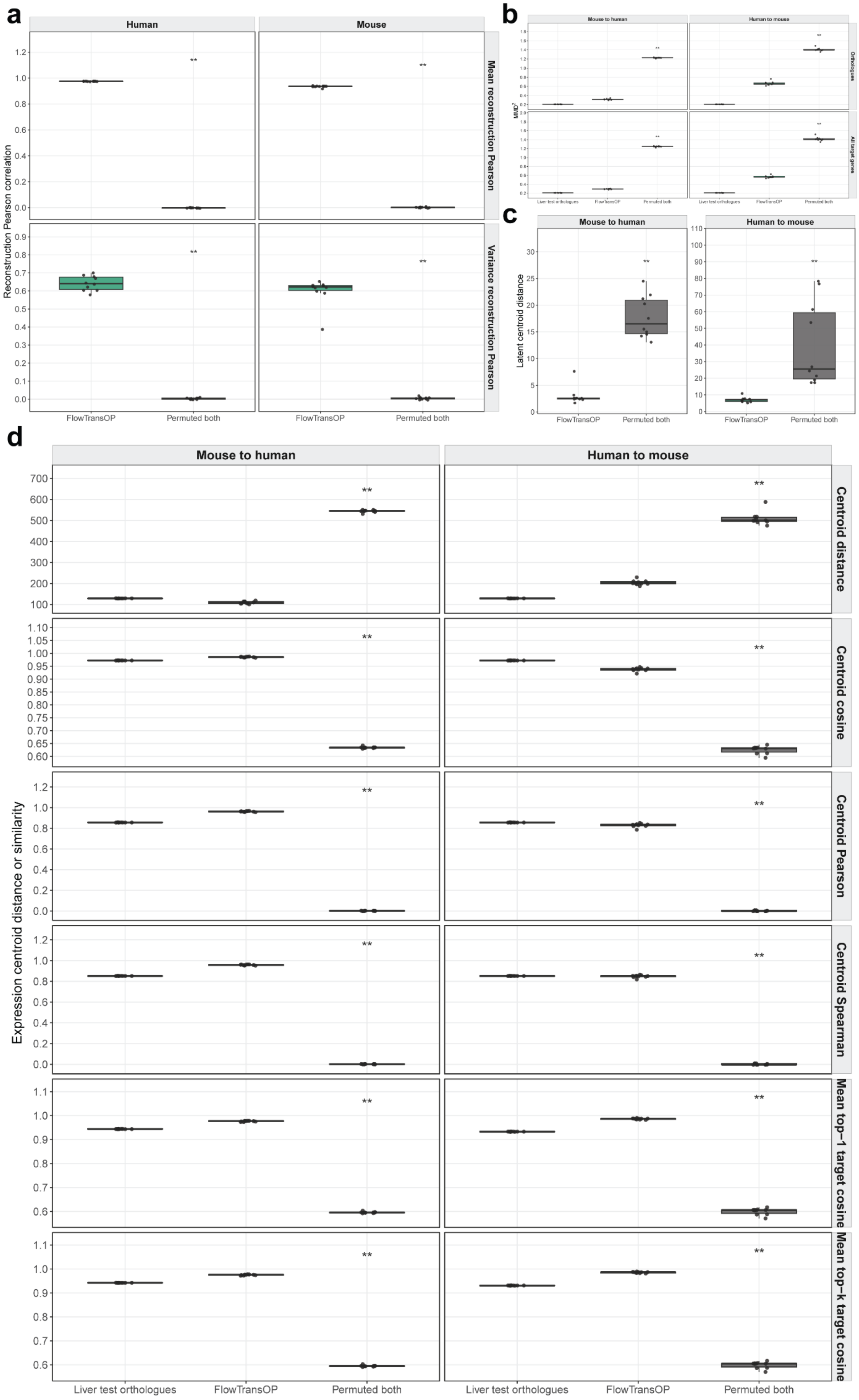
a
Human
Mouse
Reconstruction Pearson correlation
Mean reconstruction Pearson
Variance reconstruction Pearson
FlowTransOP
Permuted both
**
b
Mouse to human
Human to mouse
Orthologues
All target genes
Liver test orthologues
FlowTransOP
Permuted both
c
Mouse to human
Human to mouse
Latent centroid distance
FlowTransOP
Permuted both
**
d
Mouse to human
Human to mouse
Expression centroid distance or similarity
Centroid distance
Centroid cosine
Centroid Pearson
Centroid Spearman
Mean top−1 target cosine
Mean top−k target cosine
Liver test orthologues
FlowTransOP
Permuted both
**

**Supplementary Figure S9: Held-out liver evaluations confirm reconstruction quality and tissue-specific distributional structure. a)** Autoencoder reconstruction of held-out human and mouse liver samples, assessed by Pearson correlation for reconstructed gene means and gene variances. **b)** Liver expression-space distributional matching measured by $MMD^2$ over orthologues and all target genes; lower $MMD^2$ indicates better target-liver distributional agreement. **c)** Latent centroid specificity, measured as distance between translated source-liver samples and the real target-liver centroid relative to the permuted baselines. **d)** Expression-space centroid and nearest-neighbor liver specificity metrics, including centroid distance, centroid cosine similarity, centroid Pearson and Spearman similarity, and mean top-k target cosine similarity. For a-d, p-values were computed using paired one-sided Wilcoxon signed-rank tests across folds comparing FlowTransOP with the permuted baselines, using the direction of the alternative hypothesis appropriate to each metric: higher for correlation/similarity metrics and lower for distance or $MMD^2$ metrics. Asterisks denote Holm-adjusted p-values: *p <= 0.05, **p <= 0.01, ***p <= 0.001. In all boxplots, the centerline denotes the median, the box denotes the 1st and 3rd quartiles, and whiskers denote points within 1.5 x IQR.

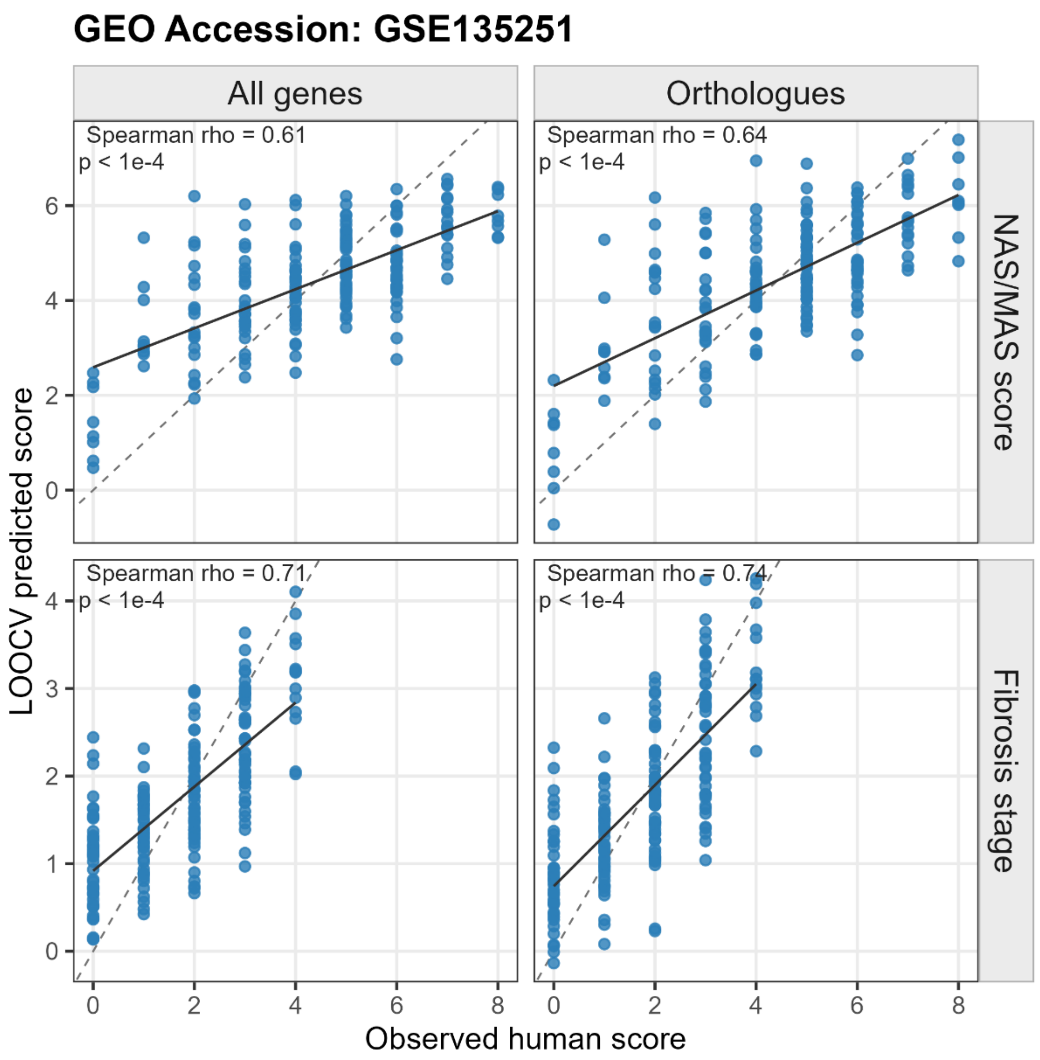


**Supplementary Figure S10: Human MASH PLSR models achieve strong leave-one-out cross-validation performance.** Partial least-squares regression (PLSR) models were trained on human Govaere et al. liver transcriptomes (GSE135251) to predict NAS/MAS score and fibrosis stage. Scatter plots compare observed clinical scores with leave-one-out cross-validated predicted scores for models trained using all genes or orthologue-restricted features. Dashed lines denote the identity line and solid lines denote the fitted trend. Spearman rho and p-values were computed by Spearman rank-correlation tests between observed and cross-validated predicted scores. These PLSR models were used to score raw mouse orthologue profiles and FlowTransOP-translated human-space profiles in the MASH case studies.

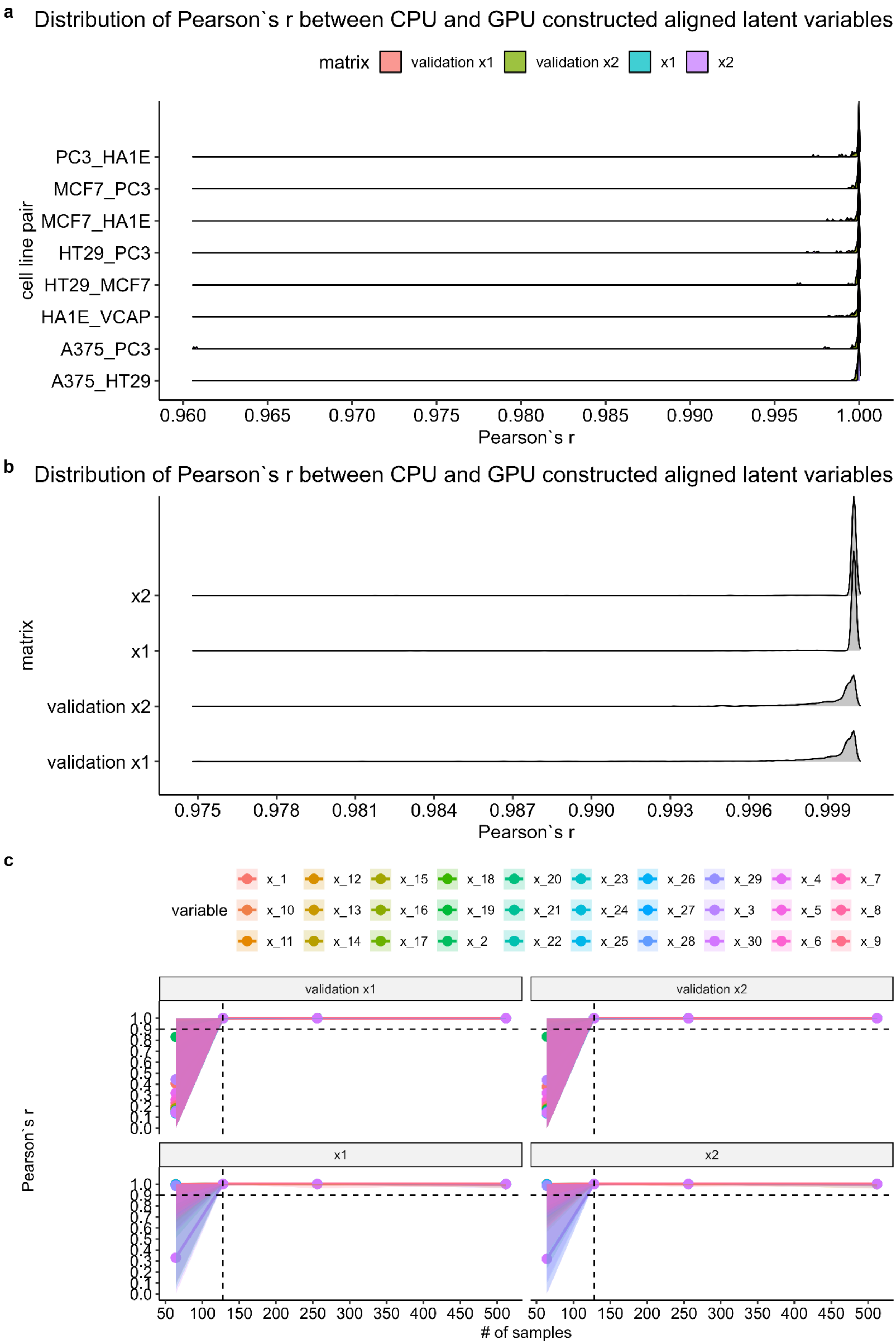
a
Distribution of Pearson`s r between CPU and GPU constructed aligned latent variables
matrix
validation x1
validation x2
x1
x2
cell line pair
PC3_HA1E
MCF7_PC3
MCF7_HA1E
HT29_PC3
HT29_MCF7
HA1E_VCAP
A375_PC3
A375_HT29
0.960 0.965 0.970 0.975 0.980 0.985 0.990 0.995 1.000
Pearson`s r
b
Distribution of Pearson`s r between CPU and GPU constructed aligned latent variables
matrix
x2
x1
validation x2
validation x1
0.975 0.978 0.981 0.984 0.987 0.990 0.993 0.996 0.999
Pearson`s r
c
variable
x_1 x_12 x_15 x_18 x_20 x_23 x_26 x_29 x_4 x_7
x_10 x_13 x_16 x_19 x_21 x_24 x_27 x_3 x_5 x_8
x_11 x_14 x_17 x_2 x_22 x_25 x_28 x_30 x_6 x_9
validation x1
validation x2
x1
x2
Pearson`s r
1.0 0.9 0.8 0.7 0.6 0.5 0.4 0.3 0.2 0.1 0.0
50 100 150 200 250 300 350 400 450 500
### of samples

**<u>Supplementary Figure S11:</u> GPU and CPU implementations of TRANSACT produce concordant pre-aligned spaces in real L1000 workflows. a)** Ridge-density distributions of Pearson correlation (r) between CPU- and GPU-generated aligned latent variables for L1000 cell-line-pair tasks, shown separately for training and validation matrices from both domains. **b)** Corresponding CPU-GPU agreement for same-cell imputed L1000 matrices. **c)** CPU-GPU agreement under random subsampling of cell-line-pair data as a function of sample size. In c, points and lines denote the mean r across iterations and shaded ribbons denote the observed minimum-to-maximum range across iterations. Values near one indicate near-identical CPU and GPU pre-alignments.

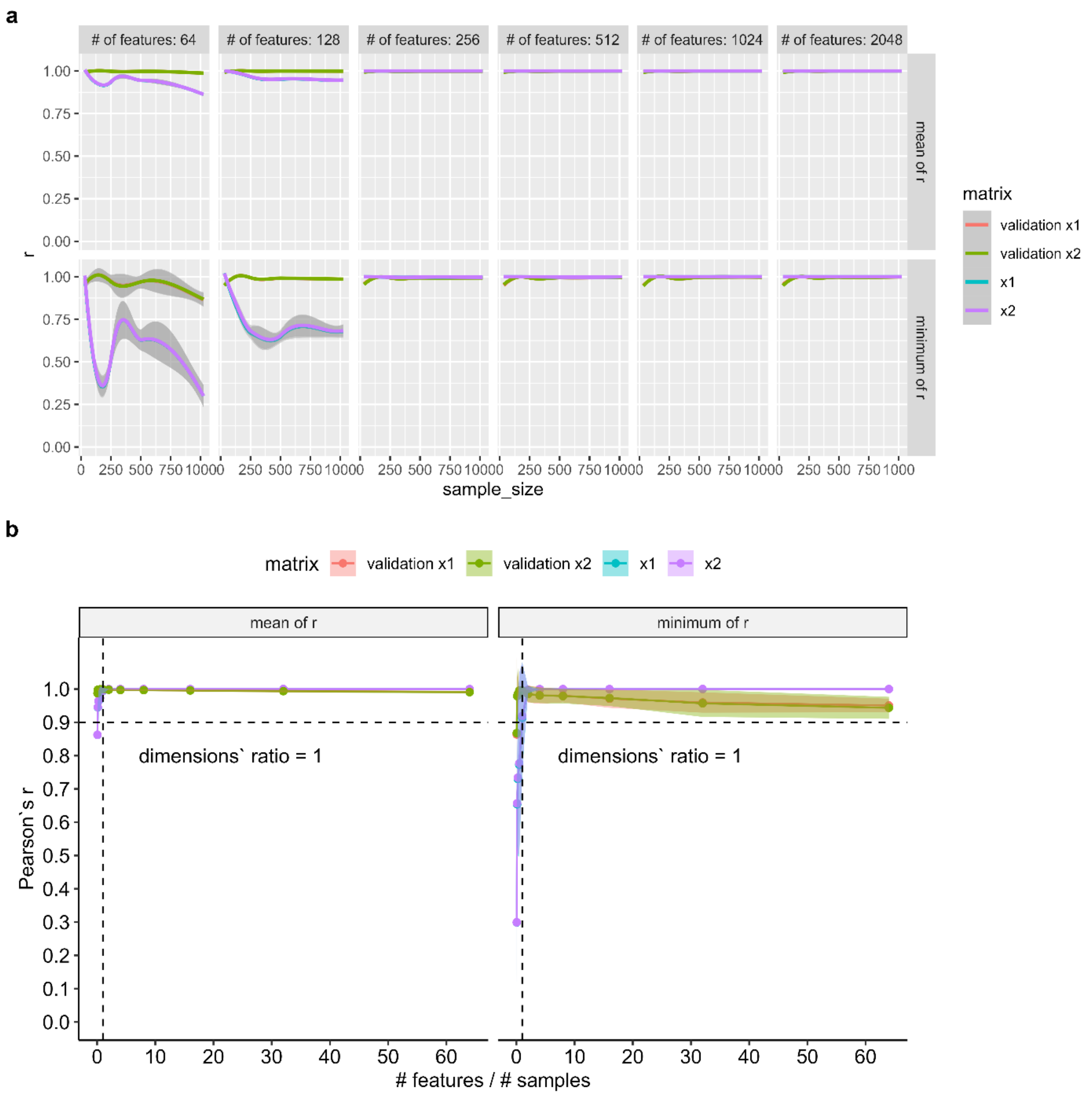


**<u>Supplementary Figure S12:</u> GPU and CPU TRANSACT implementations remain concordant for simulated data drawn from a normal distribution. a)** Pearson correlation (r) between CPU- and GPU-generated aligned latent variables as a function of sample size, faceted by feature dimension and summarized by mean r and minimum r across aligned variables. Separate curves are shown for source, target, and validation matrices. Shaded bands around smooth curves denote the default 95% confidence interval of the loess smooth. **b)** Mean r and minimum r summarized as a function of the feature-to-sample ratio. Points and lines denote the mean across iterations, shaded ribbons denote plus/minus one standard deviation (SD), the horizontal dashed line marks r = 0.9, and the vertical dashed line marks a feature-to-sample ratio of one.

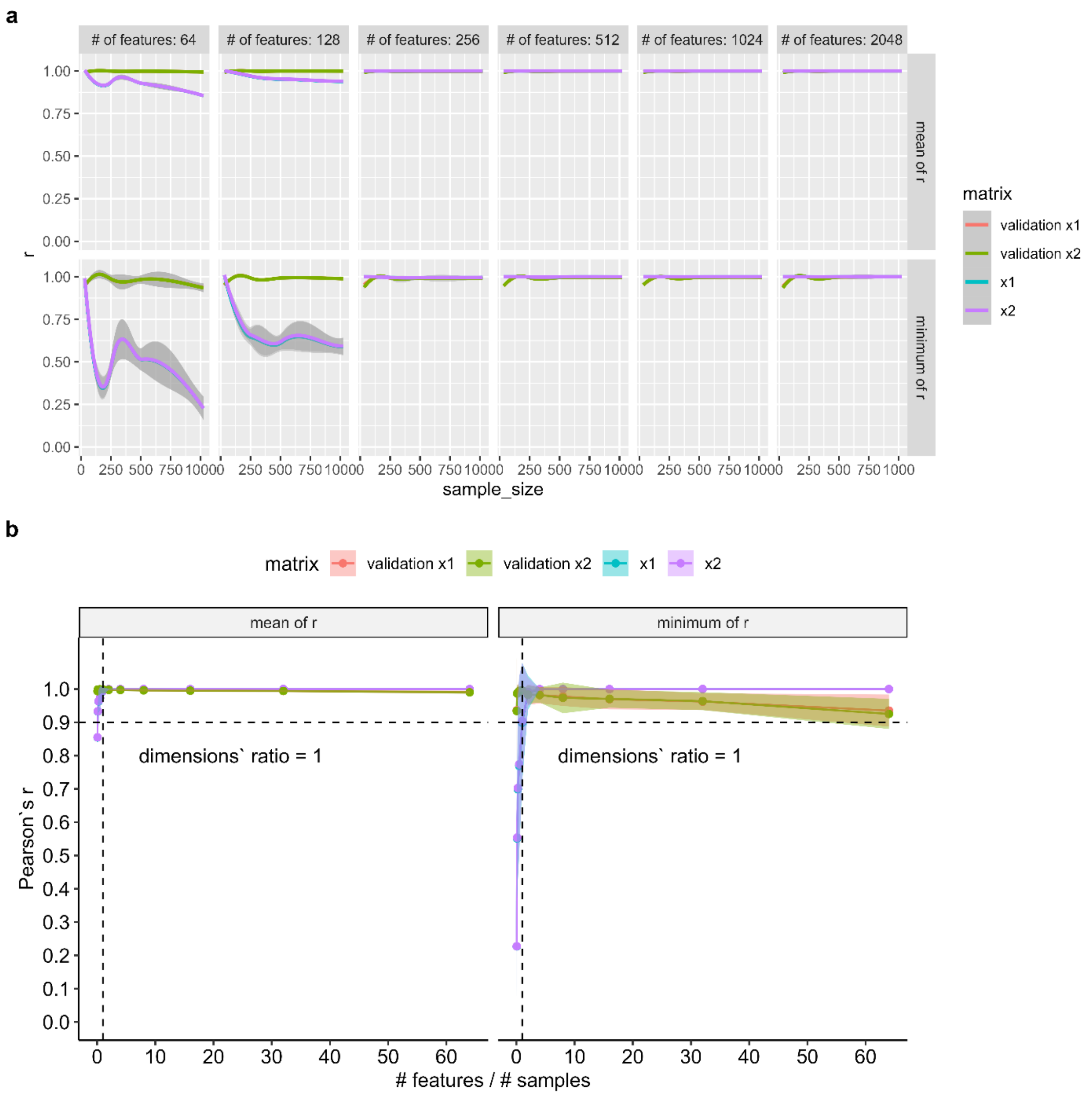


**Supplementary Figure S13: GPU and CPU TRANSACT implementations remain concordant for simulated data drawn from an exponential distribution. a)** Pearson correlation (r) between CPU- and GPU-generated aligned latent variables as a function of sample size, faceted by feature dimension and summarized by mean r and minimum r across aligned variables. Separate curves are shown for source, target, and validation matrices. Shaded bands around smooth curves denote the default 95% confidence interval of the loess smooth. **b)** Mean r and minimum r summarized as a function of the feature-to-sample ratio. Points and lines denote the mean across iterations, shaded ribbons denote plus/minus one SD, the horizontal dashed line marks r = 0.9, and the vertical dashed line marks a feature-to-sample ratio of one.

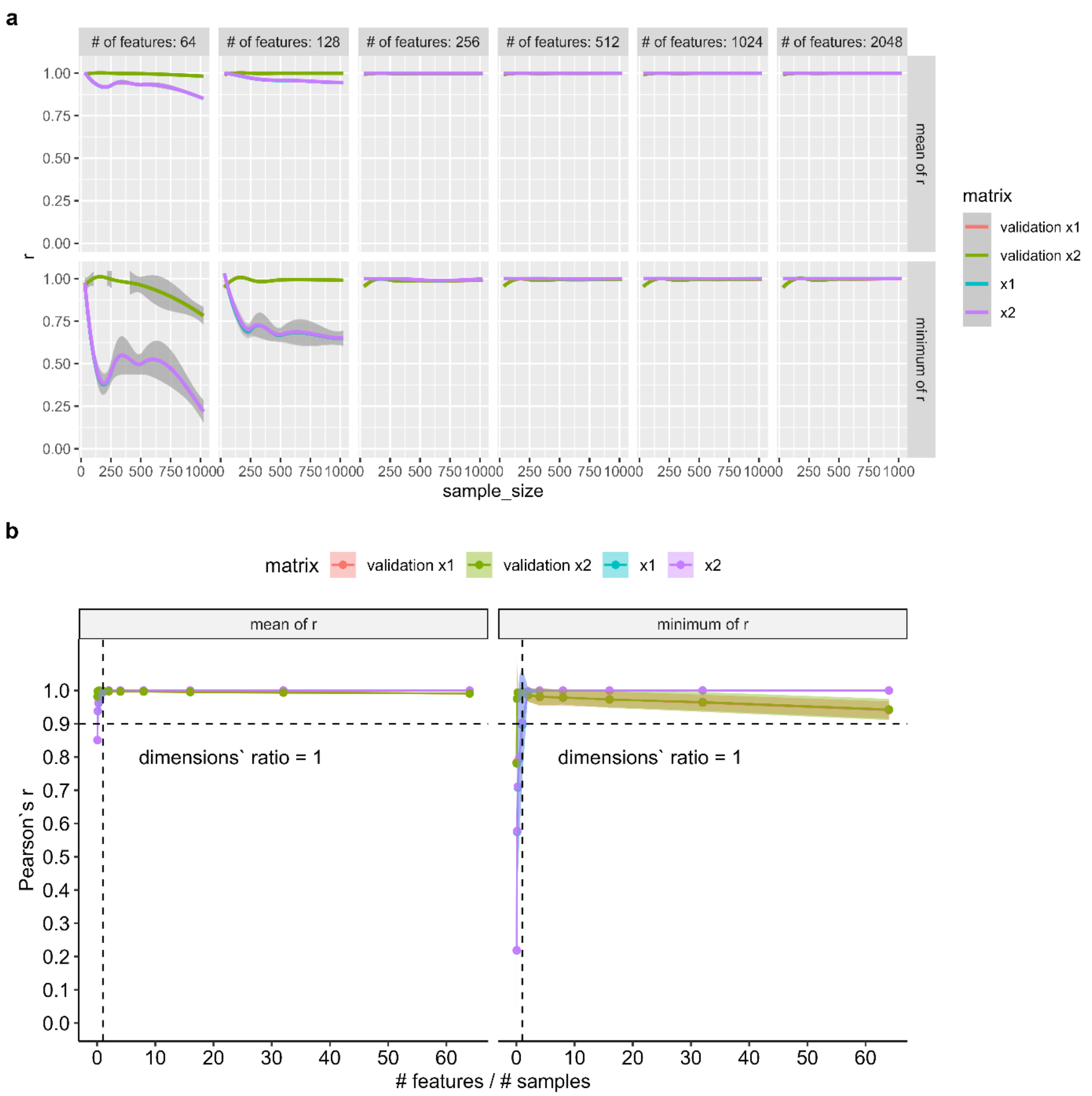


**<u>Supplementary Figure S14:</u> GPU and CPU TRANSACT implementations remain concordant for simulated data drawn from a gamma distribution. a)** Pearson correlation (r) between CPU- and GPU-generated aligned latent variables as a function of sample size, faceted by feature dimension and summarized by mean r and minimum r across aligned variables. Separate curves are shown for source, target, and validation matrices. Shaded bands around smooth curves denote the default 95% confidence interval of the loess smooth. **b)** Mean r and minimum r summarized as a function of the feature-to-sample ratio. Points and lines denote the mean across iterations, shaded ribbons denote plus/minus one SD, the horizontal dashed line marks r = 0.9, and the vertical dashed line marks a feature-to-sample ratio of one.

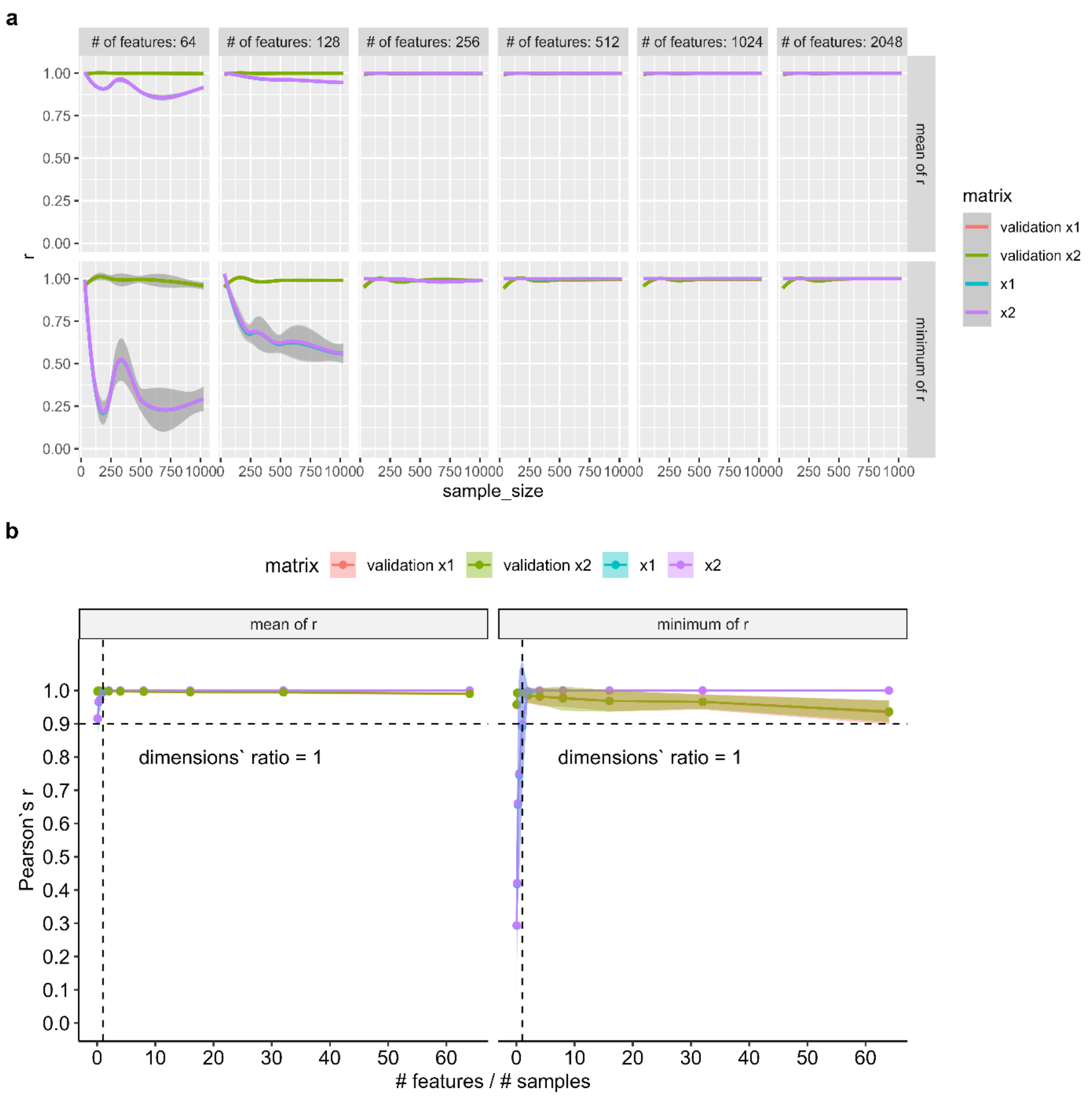


**<u>Supplementary Figure S15:</u> GPU and CPU TRANSACT implementations remain concordant for simulated data drawn from a log-normal distribution. a)** Pearson correlation (r) between CPU- and GPU-generated aligned latent variables as a function of sample size, faceted by feature dimension and summarized by mean r and minimum r across aligned variables. Separate curves are shown for source, target, and validation matrices. Shaded bands around smooth curves denote the default 95% confidence interval of the loess smooth. **b)** Mean r and minimum r summarized as a function of the feature-to-sample ratio. Points and lines denote the mean across iterations, shaded ribbons denote plus/minus one SD, the horizontal dashed line marks r = 0.9, and the vertical dashed line marks a feature-to-sample ratio of one.

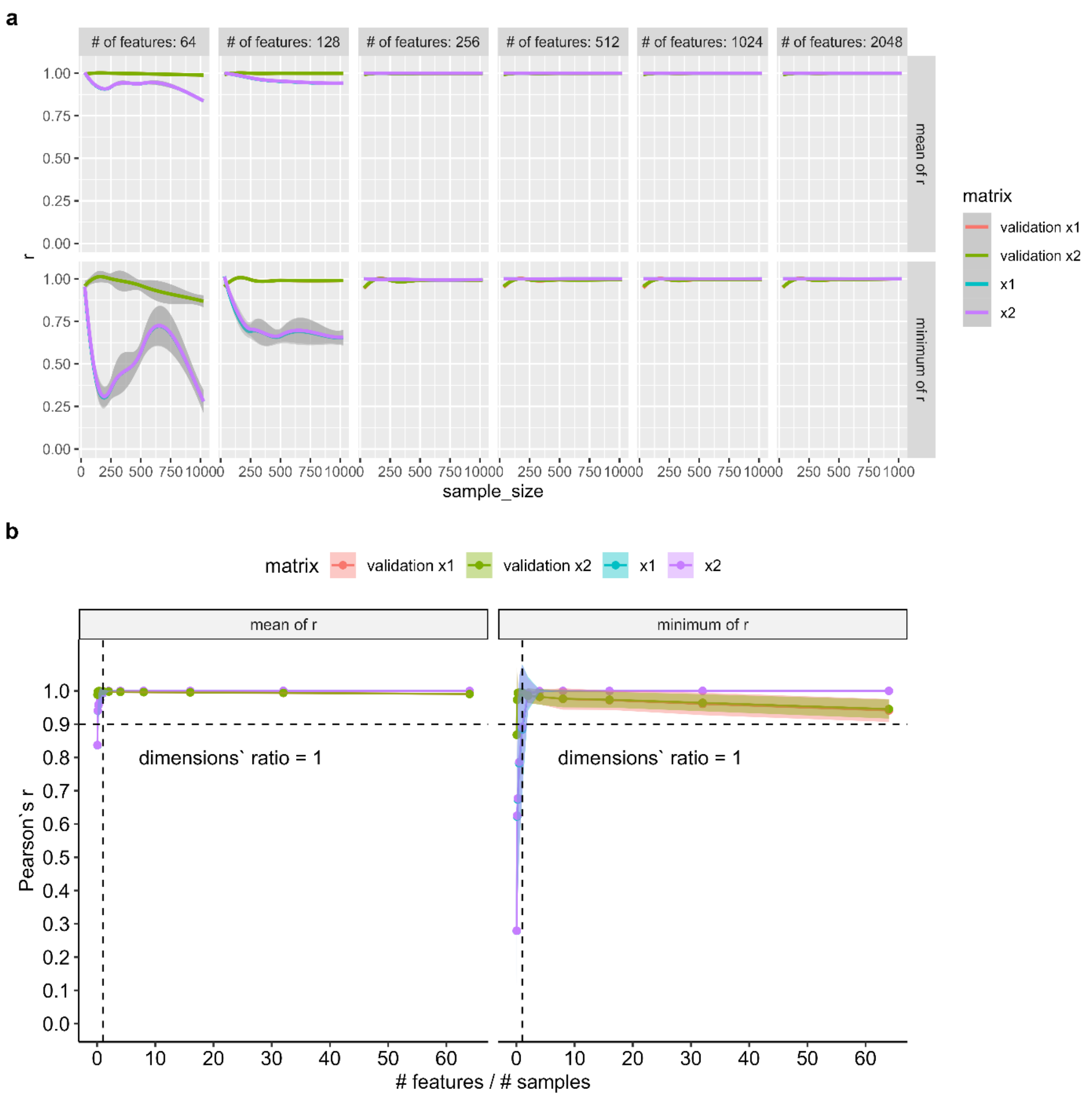


**<u>Supplementary Figure S16:</u> GPU and CPU TRANSACT implementations remain concordant for simulated data drawn from a Dirichlet distribution. a)** Pearson correlation (r) between CPU- and GPU-generated aligned latent variables as a function of sample size, faceted by feature dimension and summarized by mean r and minimum r across aligned variables. Separate curves are shown for source, target, and validation matrices. Shaded bands around smooth curves denote the default 95% confidence interval of the loess smooth. **b)** Mean r and minimum r summarized as a function of the feature-to-sample ratio. Points and lines denote the mean across iterations, shaded ribbons denote plus/minus one SD, the horizontal dashed line marks r = 0.9, and the vertical dashed line marks a feature-to-sample ratio of one.

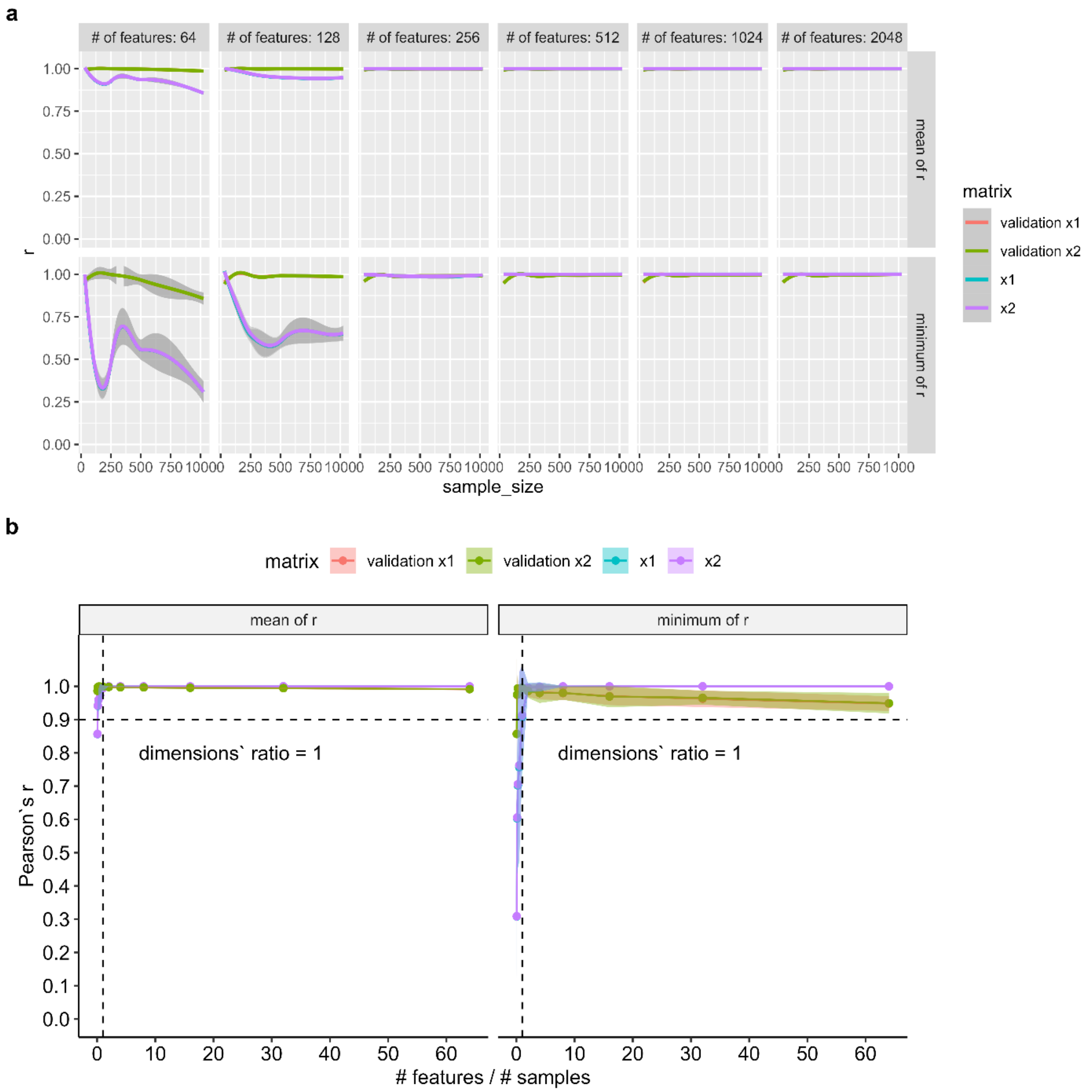


**<u>Supplementary Figure S17:</u> GPU and CPU TRANSACT implementations remain concordant for simulated data drawn from a Poisson distribution. a)** Pearson correlation (r) between CPU- and GPU-generated aligned latent variables as a function of sample size, faceted by feature dimension and summarized by mean r and minimum r across aligned variables. Separate curves are shown for source, target, and validation matrices. Shaded bands around smooth curves denote the default 95% confidence interval of the loess smooth. **b)** Mean r and minimum r summarized as a function of the feature-to-sample ratio. Points and lines denote the mean across iterations, shaded ribbons denote plus/minus one SD, the horizontal dashed line marks r = 0.9, and the vertical dashed line marks a feature-to-sample ratio of one.

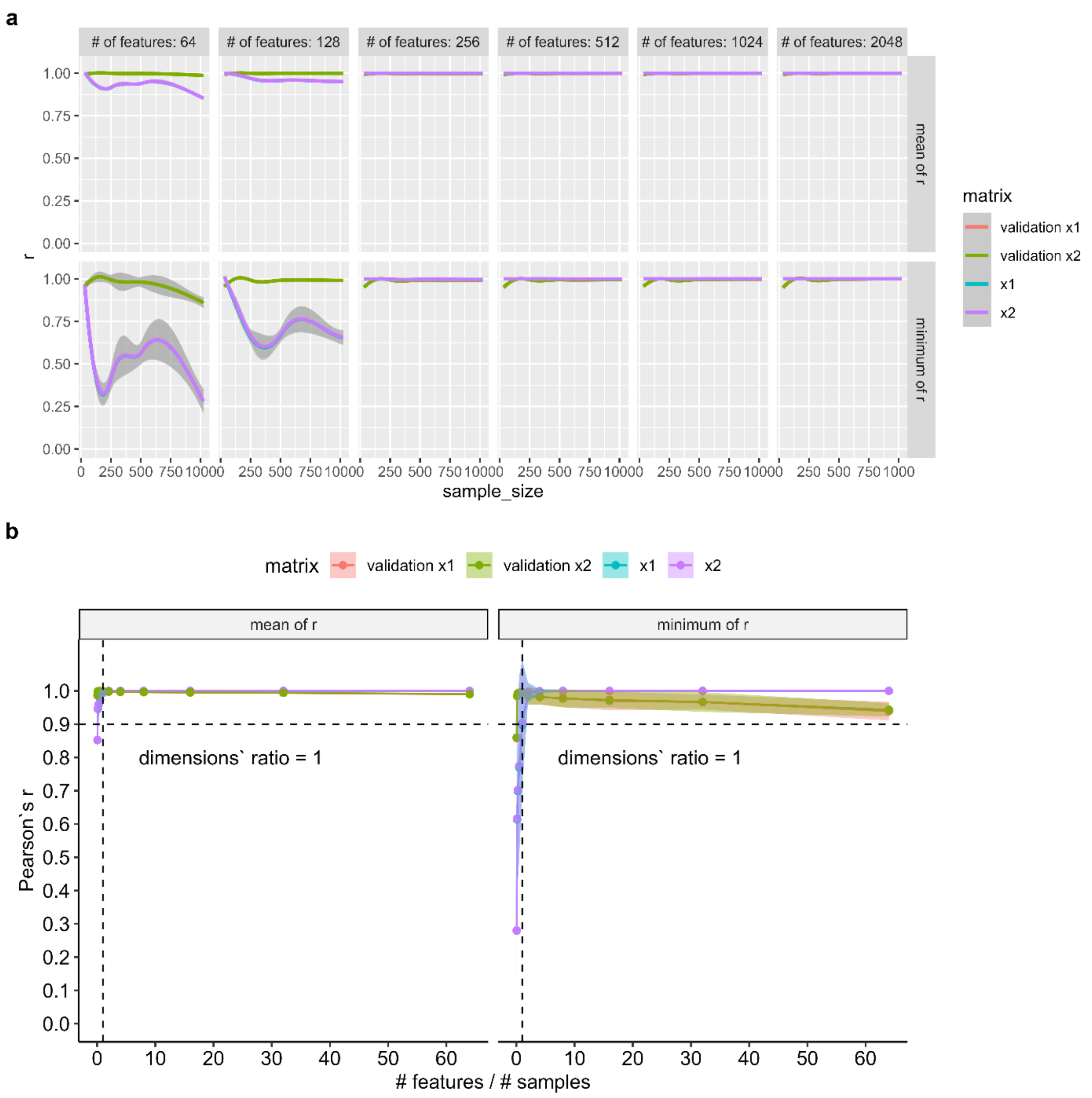


**<u>Supplementary Figure S18:</u> GPU and CPU TRANSACT implementations remain concordant for simulated data drawn from a uniform distribution. a)** Pearson correlation (r) between CPU- and GPU-generated aligned latent variables as a function of sample size, faceted by feature dimension and summarized by mean r and minimum r across aligned variables. Separate curves are shown for source, target, and validation matrices. Shaded bands around smooth curves denote the default 95% confidence interval of the loess smooth. **b)** Mean r and minimum r summarized as a function of the feature-to-sample ratio. Points and lines denote the mean across iterations, shaded ribbons denote plus/minus one SD, the horizontal dashed line marks r = 0.9, and the vertical dashed line marks a feature-to-sample ratio of one.

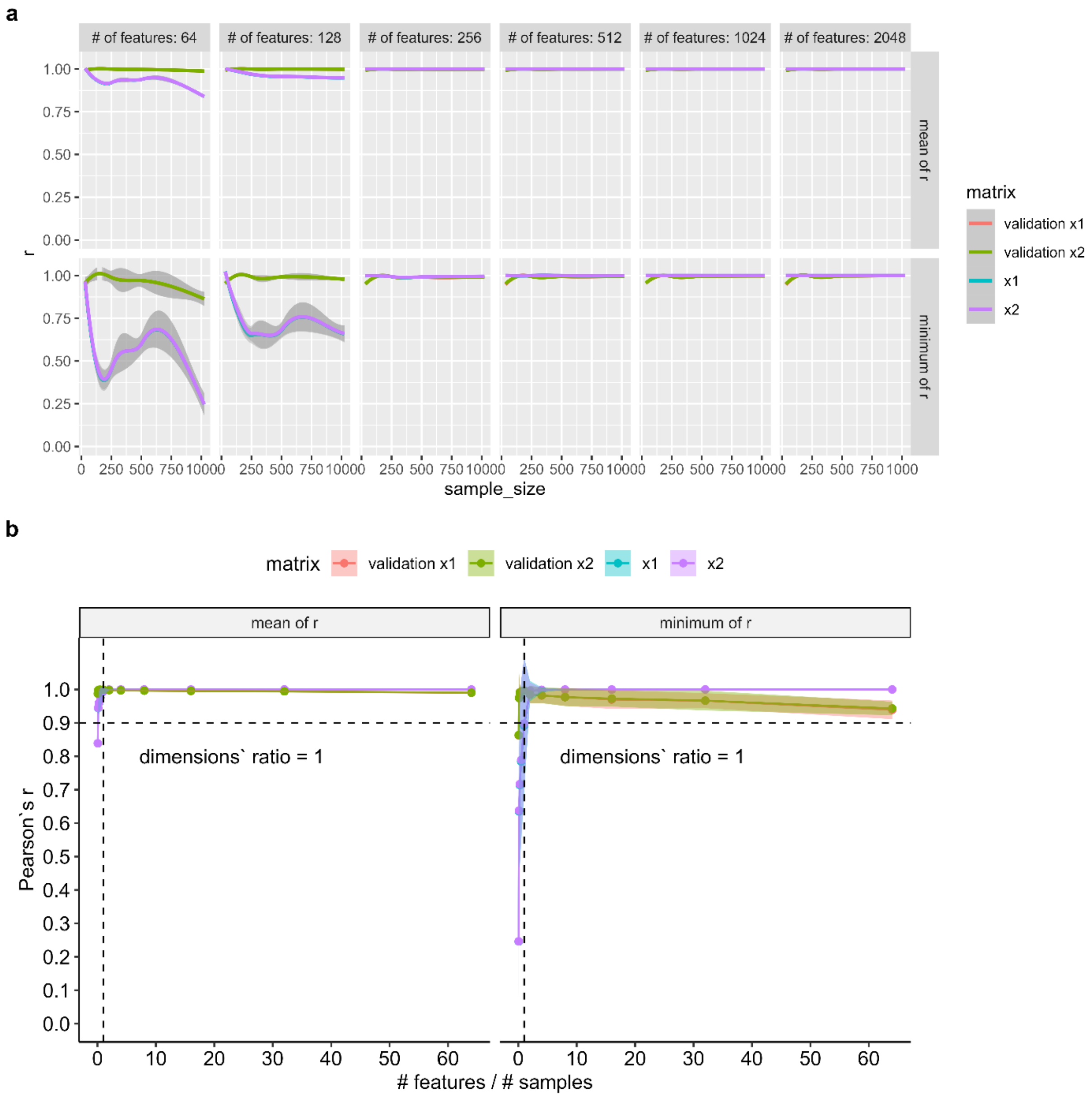


**Supplementary Figure S19: GPU and CPU TRANSACT implementations remain concordant for simulated data drawn from a Xavier-initialized distribution. a)** Pearson correlation (r) between CPU- and GPU-generated aligned latent variables as a function of sample size, faceted by feature dimension and summarized by mean r and minimum r across aligned variables. Separate curves are shown for source, target, and validation matrices. Shaded bands around smooth curves denote the default 95% confidence interval of the loess smooth. **b)** Mean r and minimum r summarized as a function of the feature-to-sample ratio. Points and lines denote the mean across iterations, shaded ribbons denote plus/minus one SD, the horizontal dashed line marks r = 0.9, and the vertical dashed line marks a feature-to-sample ratio of one.
